# Self-supervised learning for characterising histomorphological diversity and spatial RNA expression prediction across 23 human tissue types

**DOI:** 10.1101/2023.08.22.554251

**Authors:** Francesco Cisternino, Sara Ometto, Soumick Chatterjee, Edoardo Giacopuzzi, Adam P. Levine, Craig A. Glastonbury

**Author notes:** Corresponding Author: Craig A. Glastonbury.

## Abstract

As vast histological archives are digitised, there is a pressing need to be able to associate specific tissue substructures and incident pathology to disease outcomes without arduous annotation. Such automation provides an opportunity to learn fundamental biology about how tissue structure and function varies in a population. Recently, self-supervised learning has proven competitive to supervised machine learning approaches in classification, segmentation and representation learning. Here, we leverage self-supervised learning to generate histology feature representations using 1.7M images across 23 healthy tissues in 838 donors from GTEx. Using these representations, we demonstrate we can automatically segment tissues into their constituent tissue substructures and pathology proportions, and surpass the performance of conventionally used pre-trained models. We observe striking population variability in canonical tissue substructures, highlight examples of missing pathological diagnoses, incorrect assignment of target tissue and cross-tissue contamination. We demonstrate that this variability in tissue composition leads to a likely overestimation of eQTL tissue sharing and drives dramatic differential gene expression changes. We use derived tissue substructures to detect 284 tissue substructures and pathology specific eQTLs. As our derived histology representations are rich morphological descriptors of the underlying tissue, we introduce a multiple instance learning model that can predict and spatially localise individual RNA expression levels directly from histology to specific substructures and pathological features. We validate our RNA spatial predictions with matched ground truth immunohistochemistry (IHC) for several well characterised marker genes, recapitulating their known spatial specificity. Finally, we derive a gene expression spatial enrichment metric, allowing us to detect genes specifically expressed within sites of pathology (e.g. arterial calcification). Together, these results demonstrate the power of self-supervised machine learning when applied to vast histological datasets to allow researchers to pose and answer questions about tissue pathology, its spatial organisation and the interplay between morphological tissue variability and gene expression.

## Introduction

Histology is a relatively inexpensive and effective technique that is commonly used to diagnose and characterise a multitude of diseases, most notably, cancer. Classically, glass histology slides are examined by a pathologist under a microscope; however, recently, there has been considerable momentum in digitising pathology workflows, as histology slides can be quickly scanned at high resolution (40×) to generate Whole Slide Images (WSI). This digitisation provides an opportunity to leverage several advances in computer vision and machine learning (ML). Indeed, multiple ML methods have been developed, largely for malignant pathological entities, to segment specific cell types, tissue structures, diagnostic features of interest^1–3^, predict the mutation status of tumours^4^ and diagnostically classify histology tissue sections^3^. Whilst such supervised learning algorithms have proved successful, they rely on expert crafted labels and typically utilise models pre-trained on ImageNet, a large collection of natural images unlike histology^5^. Recently, self-supervision has proven to be a useful methodology to learn rich, low-dimensional representations of imaging data that have shown competitive performance to supervised methods, but can be used for a wide variety of downstream tasks, with some preliminary applications focused on histopathology^6^.

In parallel, there are ongoing large-scale research efforts to collect both histology and paired molecular data from thousands of samples, including RNA sequencing (RNA-seq) and Whole Genome Sequencing (WGS)^7,8^. Such datasets provide an opportunity to learn how tissue structure and function vary in a population, and how constituent elements of tissue, in both health and disease, are impacted by both common genetic variation and gene expression. Previous efforts have focused on supervised approaches, by extracting and quantifying the size and distribution of specific cell-types of interest and characterising them epidemiologically and genetically, not considering RNA-seq^1^. However, this requires the manual collection of binary segmentation labels of cells which is time consuming and therefore does not scale to multiple tissue types. Supervised methods utilising ImageNet pre-trained models have also been developed that aim to predict gene expression directly from histology image tiles, but have not sought to decompose gene expression contributions from underlying tissue substructures and pathological features present in the tissue^9^. Additionally such efforts have been entirely focused on malignant pathological entities, in which the histology changes are usually considerable and provide a very strong signal for the algorithm to learn from as compared to natural histological variation in healthy donors^3,10^. Seminal work on unsupervised approaches that have aimed to couple histology, gene expression and genetic variation have focused on the use of latent factor models^11–13^. These approaches have been able to characterise both shared and specific sources of gene expression and histological variation and have described “image QTLs”, in which genetic variants drive changes in tissue morphology. However, these latent factors are abstract and therefore it is not always possible to understand the underlying biological and pathological processes they are capturing^11–13^. Finally, the extent to which specific histological tissue substructures and pathological features vary naturally in a population, as quantified computationally, and hence objectively, from large numbers of WSI, has not been widely addressed, nor how such variation can be associated to common genetic variants or changes in gene expression related to tissue morphology.

We advance previous work by exploiting several recent ML innovations, namely Vision Transformers (ViT)^14^ coupled with self-supervised learning^15^ to combine histology, gene expression and common germline genetic variation in 9,068 samples, representing 23 distinct tissues, from 838 donors and a total of 1.7M images^8^ (**Figure 1**). We start by learning low-dimensional representations of histology tissue tiles using DINO, demonstrating that our representations are able to identify and cluster specific tissue substructures, cells and pathological features without labels better than previous ImageNet pre-trained models which are the standard in the field (**Figure 1a-e**).

**Figure 1:**
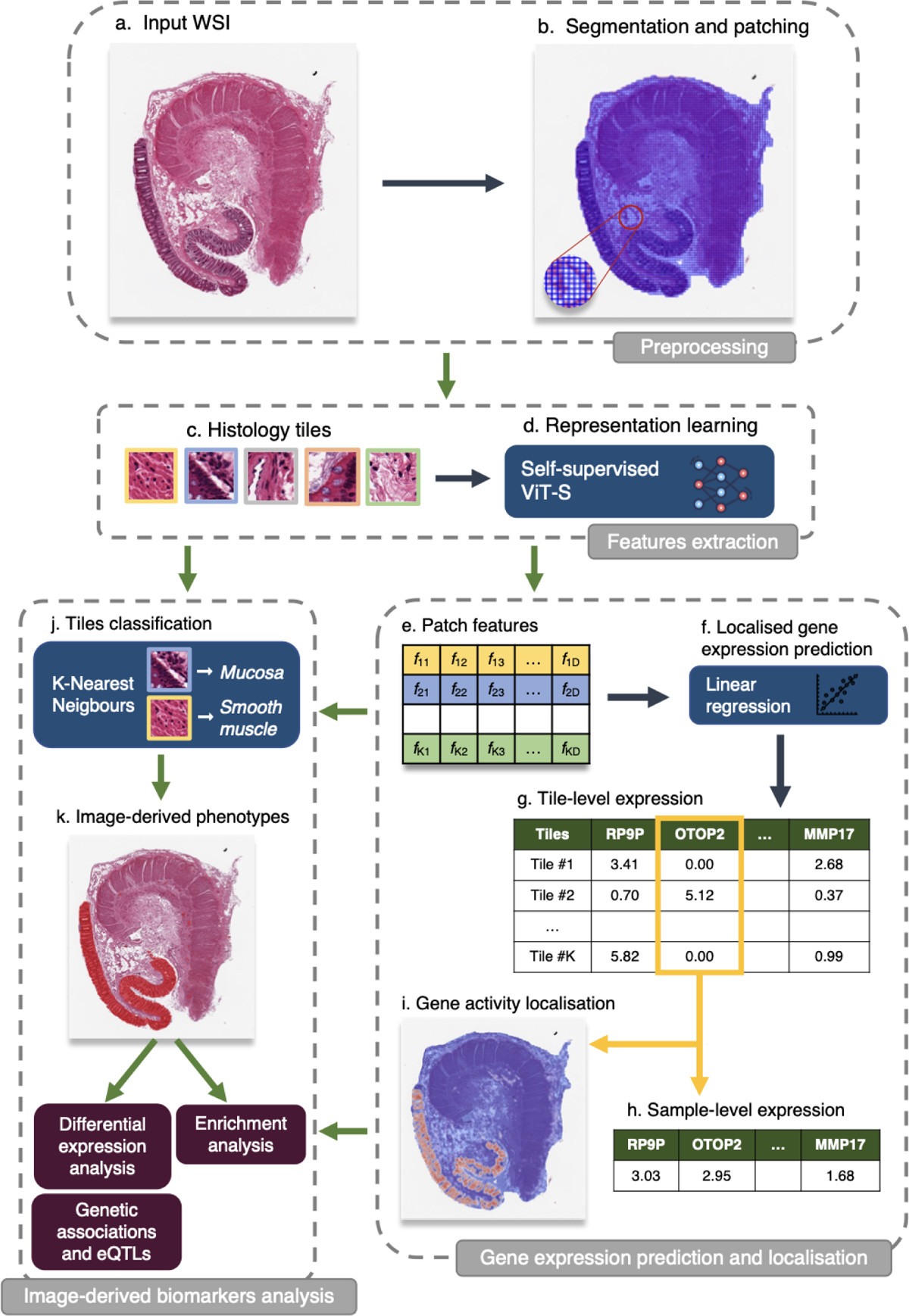
Schematic representation of using self-supervised representations learnt from whole slide image histology for segmentation of tissue substructures, pathological features and understanding morphology-expression-genetic associations using *RNAPath*.

We utilise the ability of these representations to identify substructures and pathological features to automatically segment all GTEx WSI, with limited manual labelling of specific features (< 0.5% of the data used) obtaining substructure and pathology proportions per donor sample (**Figure 1j**). Next, using these derived substructures and pathological features, we characterise profound tissue variability across donors that drives substantial differential gene expression, as well as characterise common germline genetic variants associated with specific histopathological features through both genome-wide association analysis (GWAS) and interaction eQTL analyses (**Figure 1k**). Finally, despite not being trained specifically for this task, as our representations are rich morphological descriptors of tissue histology both within and across donors, we are able to predict and spatially localise individual RNA expression levels with superior performance to competing methods (**Figure 1e-i**). We validate our spatial RNA expression predictions using ground-truth immunohistochemistry (IHC) for several canonical marker genes and subsequently characterise the specific localised expression signatures of 29 individual substructures and pathological features.

## Results

### Histology tile representations learnt via self-supervision distinguish tissue substructures and pathological features without labels

We utilised 9,068 Whole Slide Images (WSI) across 23 tissues collected from 838 donor individuals as part of GTEx^8^. WSI are gigapixel images (e.g. 50,000×150,000 pixels), in which much of the image does not contain tissue. To obtain only tissue containing sections of the WSIs, we segmented the tissue from background in each WSI using a previously published U-net (**Supplementary** Figure 1). The number of image tiles (128×128 pixels) derived from WSIs in GTEx varies dramatically, with the lowest being 248 in a tibial nerve WSI versus 82,605 in a testis WSI. This range reflects the variation in sampled tissue section sizes. As it is computationally intractable to process the collection of raw image tiles per WSI on a GPU, as with previous work, we embedded the image tiles into 348-dimensional representations (see Methods). Contrary to previous work that utilise CNN models pre-trained on ImageNet, a dataset of natural images unlike histology, we train a self-supervised model on GTEx directly^3^. Self-supervised models have been recently shown to be effective in learning compact, rich image representations^6^. Therefore, we sought to learn relevant features present in histology images by training a Vision Transformer (ViT-S) on 1.7M GTEx histology patches using the self-supervised DINO framework^15,16^ (see Methods). Despite being trained with no labels, we see that learnt representations clearly capture and separate cell types (e.g. adipocytes), pathological features (e.g. arterial calcification) and tissue substructures (e.g. tibial vessel layers: intima, media, adventitia) (Figure 2**, Supplementary** Figure 2 for additional tissues). We compared our self-supervised representations learnt from GTEx, to the commonly used representations obtained from a ResNet50 model pre-trained on ImageNet. We see that our self-supervised representations better capture intrinsic tissue substructures with a consistent improvement in maximum silhouette score across a range of k-mean clusters (0.15 vs 0.24 for esophagus mucosa and 0.17 vs 0.21 for tibial artery, for k = [3-20]) (**Supplementary** Figure 3).

**Figure 2.**
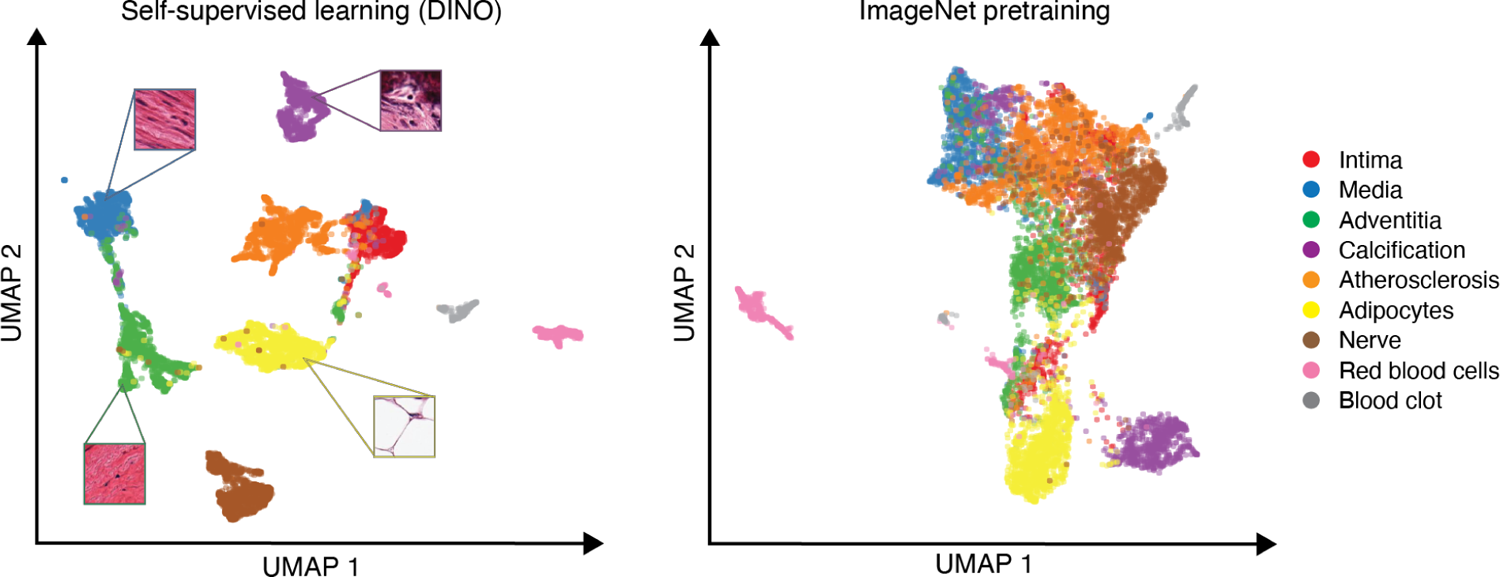
UMAP embeddings of tibial artery patch features from self-supervised ViT-S (trained using DINO) versus ResNet50 with pretrained weights from ImageNet. Patches have been manually labelled with tissue substructures/pathologies to interpret clusters.

### Self-supervised WSI tissue substructure and pathology segmentation

Segmenting various tissues into constituent tissue substructures is a time consuming task that does not scale easily to thousands of WSI. Additionally, it has not been widely documented (across thousands of samples with objective computationally derived quantification) how normal tissue substructure varies histologically in a population or what genes are specific to each substructure or pathological feature. To automate the segmentation of tissue present in WSI into constituent tissue substructures and pathological features, we manually labelled a small subset of image patches (< 0.50% tiles of overall dataset) from a subset of tissue types, with annotations that were validated by a clinical histopathologist (see Methods). Using the DINO representations obtained from these labelled image tiles, we trained a k-Nearest Neighbours (kNN) model and inferred the class of each unlabelled tile representation (see Methods). Qualitatively, we see that inferred tile labels clearly recover known tissue substructures and pathological features when overlayed with their label on their corresponding WSI (Figure 3). We quantitatively evaluated the accuracy of the kNN in the patch-level classification: for each tissue, we held-out 10% of the annotated patches of each class from the model fitting and measured its accuracy across 10 folds. Median accuracy across all derived tissue substructure and pathological features in all tissues was 92%.

**Figure 3.**
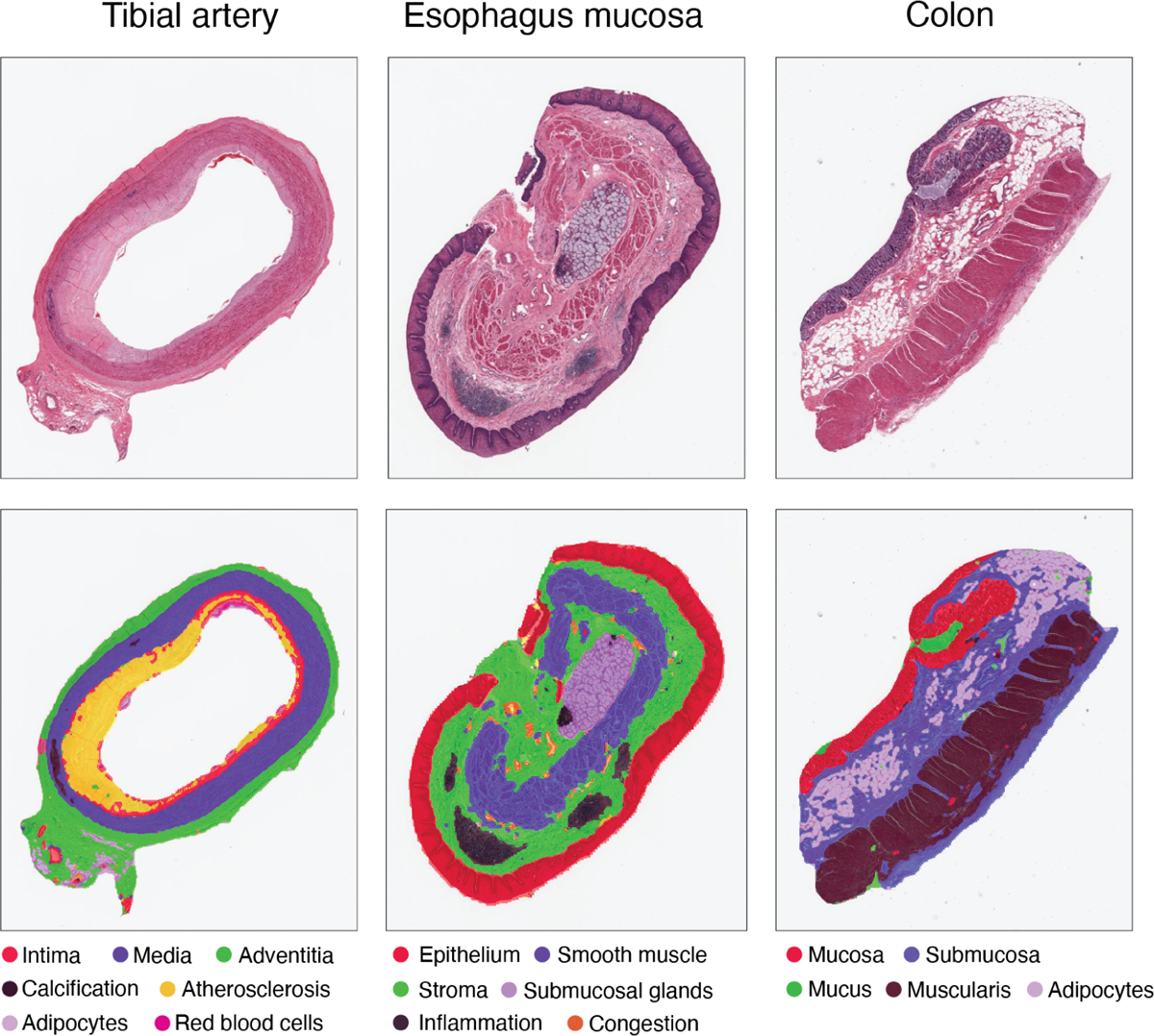
Segmentation of tissue samples from tibial artery, esophagus mucosa and colon into substructures and localised pathological features via K-Nearest neighbours on patch features. Tissue types have been labelled according to their GTEx descriptor which may not perfectly represent the specimen (e.g. the esophagus mucosa example includes submucosa).

To further assess our automated segmentation quantitatively and also evaluate the gain in information of our tissue substructure and pathological features, we used cases in which samples had been described as calcified in the GTEx pathology notes versus our inferred calcification labelled tiles per WSI. Tiles inferred as belonging to the calcification class had high sensitivity in recovering ground-truth pathologist labels, with AUC = 0.94 (**Supplementary** Figure 4), indicating that the vast majority of true positive cases were correctly identified by the kNN segmentation model. Additionally, when considering calcification occupying > 5% of the tissue present in a WSI, our model identifies a further four cases. Upon manual inspection, two of these four cases were false positives containing debris that resembled calcification; however, two WSI clearly contained calcification that was not labelled as such in the GTEx pathology notes (**Supplementary** Figure 5). This highlights the utility of our approach to discover unannotated pathological features, with the potential to aid pathological reporting in vast digitised histological datasets. The full list of tissue substructures and pathological features with corresponding accuracy are present in **Supplementary Table 1**.

### Substantial variability in tissue substructure proportions across donors

Whilst GTEx pathology notes contain labels of whether subjects have a given pathological feature or not, there is no information on its extent, i.e. the proportion of affected tissue, nor its spatial location within the tissue. Using our kNN segmentation model, we inferred labels for all tissue tiles across all WSI considered. By doing so, we can represent any given sample as the proportion of its inferred tissue substructures and pathological features, allowing us to quantify inter-subject variability. We see that the proportion of different tissue substructures and pathological features vary dramatically across donors within the same tissue type (Figure 4**, Supplementary** Figure 6). For example, the proportion of calcified tissue in tibial artery samples varies from 0 to 44% with a mean of 3.3%. This pattern is true across all tissues and across substructures quantified (**Supplementary Table 2**). In some but not all cases, this likely represents tissue sampling variation as opposed to true biological or pathological variation.

**Figure 4:**
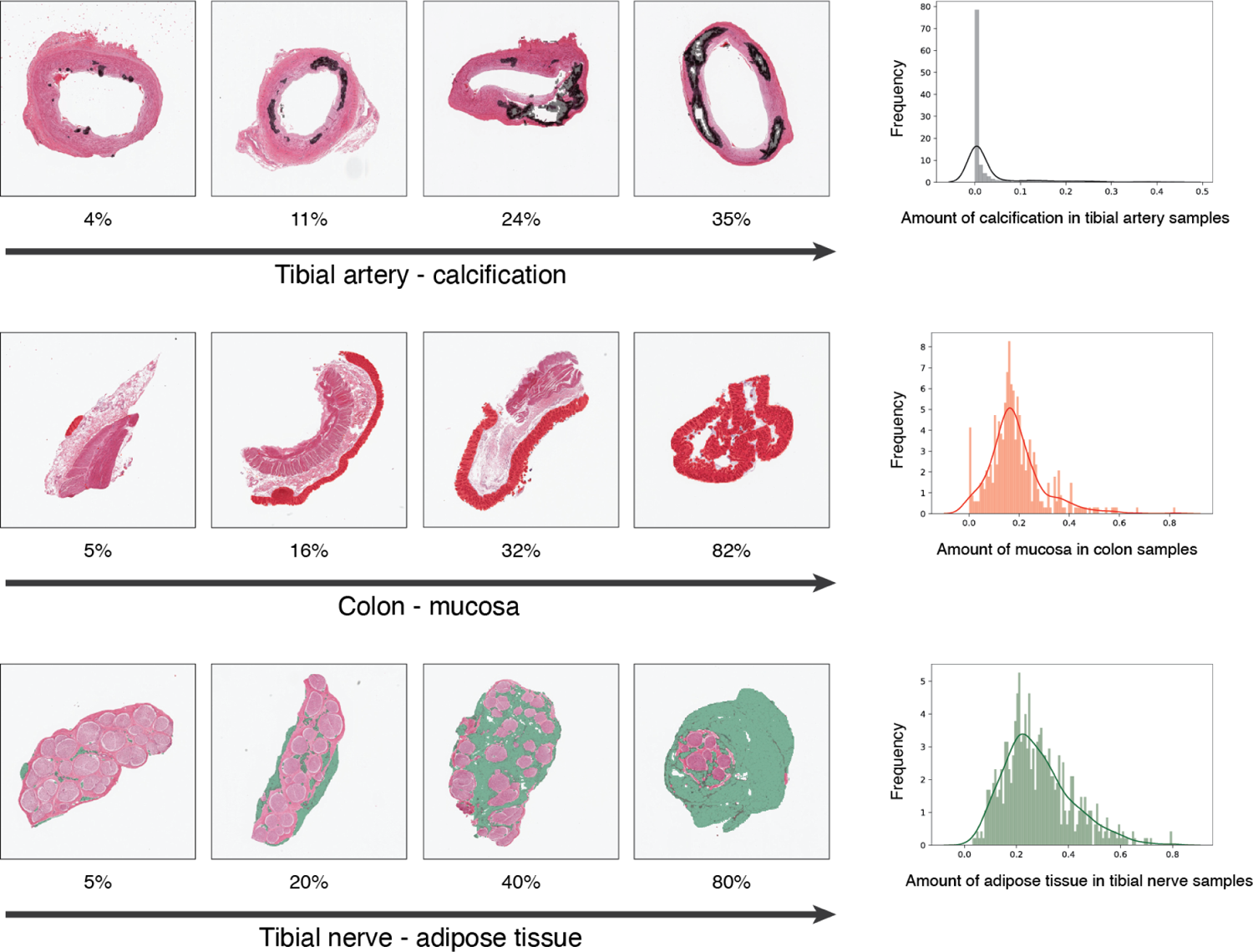
Example of two quantified tissue substructures (e.g. adipose tissue in tibial nerve and mucosa in colonic tissue) and one pathology entity (e.g. calcification in tibial artery) and their distribution. Significant variability is observed for both canonical tissue substructures and pathological features across donors.

An acute difficulty when dissecting tissue without laser capture microdissection (LCM) is obtaining the correct target tissue of interest with little to no contamination of other tissue components. This is critical for enabling the precise characterisation of gene expression tissue specificity or quantifying the degree of tissue sharing across eQTLs^17–19^. For example in GTEx, ‘esophagus mucosa’ tissue is defined as having mucosal epithelium present, whilst ‘esophagus muscularis’ tissue should not. To assess the presence of a contaminant or incorrect target tissue, we assessed the degree to which squamous epithelium was present in muscularis samples. Surprisingly, 6% of muscularis samples (total n = 950) contain mucosal tissue (>1%) (**Supplementary** Figure 7). To determine whether this is recapitulated at the level of gene expression, we assessed the expression level of a gene specific to mucosal epithelium, *KRT6A*. We see that there is substantial expression of *KRT6A* in 8.8% of GTEx esophagus muscularis samples (> 20TPM), confirming both the histology and RNA-seq contain non-target tissue.

Similarly, varying amounts of adherent adipose tissue are commonly present in a variety of GTEx samples due to imperfect histological dissection. For example, 84% of coronary arteries and 95% of tibial nerve samples have > 10% of the specimen composed of adipose tissue. Indeed, using a highly specific gene expression marker of adipocytes, *PLIN1*, we see that after subcutaneous adipose tissue (median TPM = 970), visceral adipose tissue (median TPM = 542) and breast mammary tissue (median TPM = 310), tibial nerve (median TPM = 40) and coronary artery (median TPM = 30) have the highest *PLIN1* expression across all 54 tissues. These findings suggest there is significant intra-tissue donor variability in GTEx histology and hence derived RNA-seq, but also substantial “contamination” of tissue substructure types across tissue classes in GTEx. Importantly, this affects estimates of eQTL tissue-sharing, with tibial nerve, mammary, subcutaneous and visceral adipose tissue having the largest degree of tissue-shared eQTL effects across tissues^17^. Whilst tissue sharing eQTLs between different fat compartments (i.e. subcutaneous and visceral) and breast tissue would be expected, sharing of adipose-nerve eQTL effects is most likely due to the adipocyte fraction present in GTEx nerve samples rather than any inherent or underlying biological sharing of nerve-tissue specific eQTLs with adipose tissue. Our findings suggest that estimates of tissue sharing eQTLs are likely inflated and that LCM, single-cell RNA-seq and spatial technologies at scale will likely revise these estimates downwards.

### Sex and age-specific variability in tissue substructure and pathological features

Having quantitative measures of tissue substructures allows us to assess the histological impact of age and other epidemiological variables on tissue structure and its variability in a population. To address this systematically, we investigated whether variability in any of the 29 tissue substructure or pathological features quantified with accuracy > 80% (see Methods) across donors had sex, age, BMI or ischemic time specific effects. We find 18 sex, 19 age, four BMI and 19 ischemic time significant associations (see Data & Code Availability to download full summary statistics). For example, we see that the amount of arterial calcification (P-value = 7.2 × 10^-15^, β = 0.033) and atherosclerosis (P-value = 1.52 × 10^-13^, β = 0.023) increase with age, but only atherosclerosis is more common in males (P-value = 1.4 × 10^-4^, β = −0.30). Many of the significant associations were confirmatory and expected, for example breast lobules being almost exclusive to female breast tissue (P-value = 2.5 × 10^-36^, β = 0.99), the amount of solar elastosis in sun exposed skin increases with age (P-value = 1.57 × 10^-32^, β = 0.04), and autolysed mucosa capturing ischemic time effects (P-value = 2.04 × 10^-34^, β = 0.001). Interestingly, we find a link between gynecomastoid hyperplasia and age (P-value = 2.81 × 10^-8^, β = 0.01). This is likely due to increased adiposity in older age (both sexes) and decreased testosterone production in older men^20^.

Finally, adipose tissue abundance in breast tissue is known to increase with age, and this increased adiposity is associated with risk of breast cancer^21^. We demonstrate our derived adipose proportions are associated with age in female breast mammary tissue samples (P-value = 8.5 × 10^-4^, β=1.9 × 10^-2^). This effect was robust to BMI adjustment (P-value = 3.5 × 10^-4^, β = 5.9 × 10^-2^) whilst the same effect was not observed in male donors, despite being better powered (P-value = 0.75, β = −1.68 × 10^-3^). This demonstrates the ability of our approach to find epidemiological links between tissue substructures and pathological features in WSI. The integration of WSI (generated from either archival material or through routine digital pathology workflows) with detailed electronic healthcare records could prove useful to discover additional novel, prognostic and epidemiologic associations.

### Pervasive differential expression driven by substructure and pathology variation across tissues

We sought to assess the extent to which gene expression is impacted by the quantified tissue substructure variation between donors across a given tissue. To do this, we performed differential expression analysis by fitting linear models for each gene and its association with each tissue substructure and pathological feature, adjusting for confounders (see Methods). We observe pervasive differential expression within tissues and their constituent tissue substructures, with median = 1753 number of genes (FDR1%) being differentially expressed (see Data & Code Availability to download full summary statistics). Interestingly, even for individual tissue substructures that make up the majority of a particular tissue, such as dermis in skin (1955 FDR1%), tunica media in tibial artery (12,810 FDR1%) and nerve bundles in tibial nerve (751 FDR1%), there is significant differential expression between samples. These findings highlight the tissue sampling variability present in GTEx, in which the underlying proportions of each tissue has substantial inter-donor variability. In the extremes, this can represent tissue samples within a tissue class that do not resemble the same underlying target tissue that was supposed to be acquired (**Supplementary** Figure 8).

We first investigated differential gene expression (DE) enrichment in substructures whose broad pattern of DE enrichment we have a priori knowledge about, as positive controls. For example, 1,484 differentially expressed genes (FDR1%) were detected for adipocyte proportion across coronary artery samples. Reassuringly, the top differentially expressed gene was *LIPE* (β = 0.46, *P-value* = 4.43 × 10^-9^), a selective marker for adipocytes, as well as *ADIPOQ*, *PLIN1*, *PLIN5* and *CIDEC* (*P-value* < 1 × 10^-7^), all genes known to be selectively expressed in adipocytes. As expected, Gene Set Enrichment Analysis (GSEA) confirmed adipose tissue to be the most likely tissue type (*P-value =* 2.33 × 10^-71^). Similar results were obtained for other well-described tissue substructures, such as submucosal glands in esophagus mucosa being enriched for genes associated with gastric epithelial cells (*P-value* = 1.39 × 10^-33^) with top DE genes including *SPDEF* (*P-value* = 9.87 × 10^-23^), a gene required for mucous cell differentiation^22^, as well as *MUC5B* (*P-value* = 6.07 × 10^-16^), a specific marker of mucin secreting epithelial cells^23^. Similarly, levels of inflammation in esophagus mucosa were enriched for peripheral blood cells (*P-value* = 8.68 × 10^-30^) with top DE genes representing broad lymphocyte markers (e.g. *LTB*, *CD5*, *CD6*, *CD48; P-values all < 1 × 10^-^*^18^). All these confirmatory results provide reassurance that our derived proportions capture specific tissue substructures and that we are able to relate inter-donor variation in such substructures to changes in RNA levels.

Similar to quantified tissue substructures, we investigated genes differentially expressed due to differential amounts of quantified pathological features across donors. For atherosclerosis proportion in tibial artery, 6,121 DE genes were detected at FDR 1%. Top cell-type enrichments were T-memory cells, NK-cells and endothelial cells (*P-value* < 1 × 10^-16^). Macrophages, known as foam cells in atherosclerotic plaques, were also enriched but to a lesser extent (*P-value =* 1.36 × 10^-3^). Finally, DE genes for atherosclerosis were also enriched for genes implicated through GWAS (nearest gene) of Moyamoya disease (18/24 genes; *P-value* < 2.0 × 10^-3^), a cerebrovascular atherosclerotic disorder^24^. Collectively, these enrichments represent the known interplay in atherosclerosis between intima endothelial cells and the chronic inflammation and fat deposition taking place in atherosclerotic arteries and suggest a common mechanistic link with cerebrovascular calcification.

For tibial artery calcification, a co-morbid pathology of atherosclerosis, we identified 1,794 differentially expressed genes (FDR1%). Two of the most significant genes were *DUSP4* (β = 0.25, *P-value* = 2.19 × 10^-16^), known to play a role in calcium homeostasis and *KCNN4* (β = 0.21, *P-value* = 2.09 × 10^-11^), a calcium activated potassium channel shown to induce vascular calcification^25^. Enrichment analysis demonstrated macrophages (P-value = 8.47 × 10^-40^) to be the most enriched cell-type. Macrophages are known to play an important role both in atherosclerosis and concurrent arterial calcification, with recruitment of macrophages shown to drive increased osteogenic calcification and display a pro-inflammatory phenotype. Given calcification is reported in the GTEx pathology notes, we sought to compare our continuous measure of calcification derived from the WSI with the reported presence or absence of calcification in the GTEx pathology notes. To do so, we divided samples (n=579) between healthy (n = 442) and calcified (n = 137) according to the pathology notes and tested for differential expression in a linear model, whilst correcting for confounders (see Methods). We identified 1,025 differentially expressed genes after FDR1% correction versus 1,794 when using our WSI-derived continuous measure of calcification. Whilst 78% of these differentially expressed genes are shared between both analyses, our results suggest we benefit from increased power when assessing continuous measures of calcification rather than just its presence or absence, as well as the identification of genes associated with amount of calcification rather than merely its presence. In summary, these DE results demonstrate we can dissect the contribution of individual tissue substructures and pathological features on gene expression variation in bulk, by using our learned histological representations to segment tissue components and gene expression variation across donors.

### Genetic association and detection of interaction eQTLs driven by tissue substructure and pathological variation

As pathologies such as calcification are complex traits, we assessed whether derived pathological feature proportions are associated with common (minor allele frequency (MAF) >5%) genetic variation. To do this, we performed GWAS on four derived pathologies: Coronary and tibial artery calcification as well as inflammation and vascular congestion in esophagus mucosa. Whilst no variants were genome-wide significant, considering suggestive hits (P-value < 1.0 × 10^-6^), we find four variants associated with four pathological features. All variants have either been previously described in relevant complex disease GWAS or are associated with relevant traits through Phenome-Wide Association Studies (PheWAS). rs971292786-C (β = 0.51, *P-value* = 1.9 × 10^-7^) is associated with levels of calcification in coronary arteries and in a FinnGen PheWAS, rs971292786-C is associated with coronary angioplasty (β = 0.055, *P-value* = 4.10 × 10^-5^), with a consistent direction of effect. Coronary angioplasty is the primary surgical procedure used to treat atherosclerotic arteries. For inflammation in esophagus mucosa, we find two variants rs111402007-A (β = 0.53, *P-value* = 7.64 × 10^-7^) and rs35779991-C (β = 0.25, *P-value* = 8.87 × 10^-7^). rs35779991-C is genome-wide significant in a GWAS for Body Mass Index (BMI) (β = 0.018, *P-value* = 9.97 × 10^-12^) whilst rs111402007-A has been previously associated with increased White Blood Cell Count (β = 0.023, *P-value* = 5.8 × 10^-7^). Effect directions are consistent with the known relationship between low-grade systemic inflammation in obesity, and WBC count. Finally, we find a single locus rs4364259-A (β = −0.19, *P-value* = 3.47 × 10^-7^) associated with vascular congestion in esophagus mucosa which has been previously associated with hydroxyvitamin-D levels (β = 0.0158, *P-value* = 2.2 × 10^-308^). Whilst these variants seemingly make sense in the light of previous GWAS results, caution is warranted until further large-scale studies which have both genome-wide common genetic data and histology are available to enable suitable replication of our findings. (**Supplementary** Figure 9).

Similar to previous efforts^11,13,26–28^, we carried out interaction eQTL analyses to identify *cis*-eQTLs whose effect is driven by the amount of tissue substructures and pathological features across donors. By fitting linear models with tissue substructure or pathological feature as an interaction term, we identified 284 interaction eQTLs (FDR 10%) in 250 unique genes across 31 different phenotypes in eight tissues for which annotations were available. Examples of such interaction eQTLs are visualised in **Supplementary** Figure 10. These analyses compare favourably to similar work in which sparse factor models were used to discover 68 abstract image morphology QTLs (imQTLs) across 8 GTEx tissues (FDR 10%)^11^, that could not be linked directly to tissue substructure or function. These results provide further evidence that many bulk-tissue eQTLs could be due to differential amounts of tissue substructure within a tissue type due to experimental sampling variation and/or due to variability and presence of pathological features across donor tissues.

### *RNAPath* accurately predicts and spatially localises genes in histology WSI

As our self-supervised histology tile representations accurately separate known tissue substructures and pathological features, and given paired RNA-seq profiles are available for each donor from the same RNAlater aliquot, we sought to assess whether gene expression influenced by specific histomorphological features could be predicted directly from H&E histology. To do so, we introduce *RNAPath,* a multiple instance learning (MIL) model that takes as input histology tile representations and outputs both spatial expression maps for each gene as well as their whole-tissue expression prediction (see Methods). To assess *RNAPath’*s ability to predict individual RNA abundance at the bulk level, we evaluated the accuracy of bulk RNA-seq prediction by measuring the Pearson correlation coefficient (r score) between predicted expression and ground truth (see Methods). Median r-score across all genes varies substantially across tissues, with the best performance in heart (median r = 0.65) and worst performance in pituitary gland (median r = 0.13) (**Supplementary Table 3**). At the individual gene level, we find that 1,468 genes can be predicted extremely accurately from histology alone, with an r-score ≥ 0.8 in at least one tissue. Whilst our histology tile representations were not learnt with the express intent of predicting RNA levels, we demonstrate superior performance against a leading deep learning method, HE2RNA^9^, across the majority of tissues analysed (+0.20 mean r-score) (**Supplementary** Figure 11). Finally, we evaluated the tissue specificity and tissue sharing nature of genes that *RNAPath* was able to regress well (r > 0.5). We see that the majority of these genes are tissue-specific, meaning that they are regressed accurately in a single tissue. However, many genes are regressed equally well in multiple tissues, and recapitulate known tissue relationships and anatomical proximity, such as esophagus muscularis with gastroesophageal junction (IoU = 0.35) and transverse colon with sigmoid colon (IoU = 0.33). This likely reflects shared tissue substructures and cell types between tissues (**Supplementary** Figure 12).

As well as bulk level predictions, *RNAPath* provides tile level (128×128 pixel) expression predictions which can be averaged to create spatial expression maps of any specific gene across a histology sample. To validate our spatial predictions in order to use *RNAPath* for uncovering novel tissue morphology-expression relationships, we sought to examine first the spatial expression of well known marker genes. To do this, we compared the spatial predictions of *PLIN1* (adipocytes), *DCD* (eccrine sweat glands), *CRNN* (mucosal epithelium) and *SLC6A19* (colonic mucosa) to ground truth immunohistochemistry (IHC) in matched tissues from the ProteinAtlas^29^. Surprisingly, we see high concordance between our spatially resolved RNA expression predictions and that of matched antibody staining, validating that we can use *RNAPath* to draw novel inferences between RNA expression and specific tissue morphology (Figure 5).

**Figure 5:**
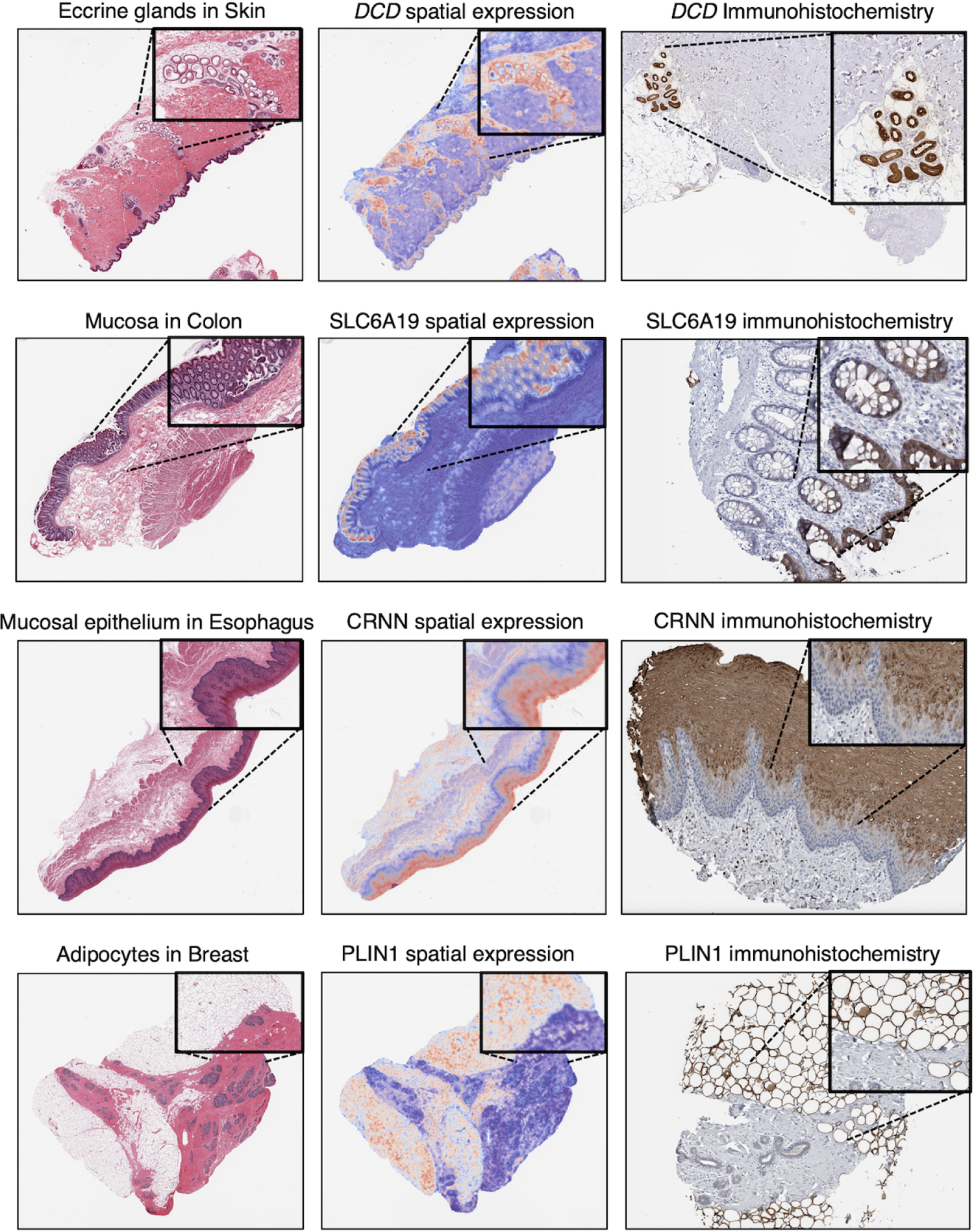
*RNAPath* predictions of canonical marker gene expression validated by immunohistochemistry (IHC). From left to right: original H&E section from GTEx, *RNAPath* predicted spatial expression for marker genes (*DCD*: Eccrine sweat glands in skin; *SLC6A19* in colonic mucosa; *CRNN* in esophageal mucosal epithelium; *PLIN1* in breast adipose tissue), IHC for corresponding protein expression in ProteinAtlas.

### Spatial expression signatures in tissue substructures and localised tissue pathological features

We sought to detail what genes were specific to individual tissue substructures and pathological features. To do this, we computed an enrichment score for every gene and every quantified tissue substructure and localised pathological feature measured. The enrichment score quantifies the difference of a gene’s predicted spatial expression between a region of interest (ROI) and the whole tissue. This measure is distinct from differential gene expression analysis. First, as our gene-wise substructure specific enrichment score (SSES) is a sample-level metric, it can be computed using a single donor (see Methods). Second, substructure enrichments are not dependent on comorbid, correlated changes within the tissue, as SSE scores are computed relative to the genes prediction across the entire histology sample. For example, a gene with high enrichment score for calcification that is also expressed in a coincident atherosclerotic plaque, will have a lower calcification enrichment score than a gene highly specific to calcification alone. This is not the case for differentially expressed genes where these two scenarios cannot be disentangled.

We investigated the relationship between up-regulated genes in our differential expression analysis (e.g. those with a positive coefficient) and our SSES metric produced by *RNAPath*. Taking as an example submucosal glands and focal inflammation in esophagus mucosa, we identified 311 and 3,471 upregulated genes respectively (FDR1%) (Figure 6). By comparing these genes to *RNAPath* SSE scores, we see that 100% and 90% have SSES > 1 for submucosal glands and inflammation respectively. For tibial artery calcification, of the 112 upregulated genes in our differential expression results, 64% have SSES > 1, highlighting the difference between differential expression induced changes and genes specific to calcification morphology (see Data & Code Availability to download full summary statistics). As lncRNAs tend to be more tissue specific we sought to test whether lncRNAs are also more tissue substructure specific than other gene biotypes. Interestingly, we observe that lncRNAs are predicted more accurately by *RNAPath* (0.35 vs 0.32 median r-score, **Supplementary** Figure 13) and are also more likely to be enriched for tissue substructures and pathologies (1.49 vs 1.23 SSES) (see Data & Code Availability to download full summary statistics). These results suggest that *RNAPath* could be used to further characterise and localise the expression of lncRNAs with unknown function.

**Figure 6:**
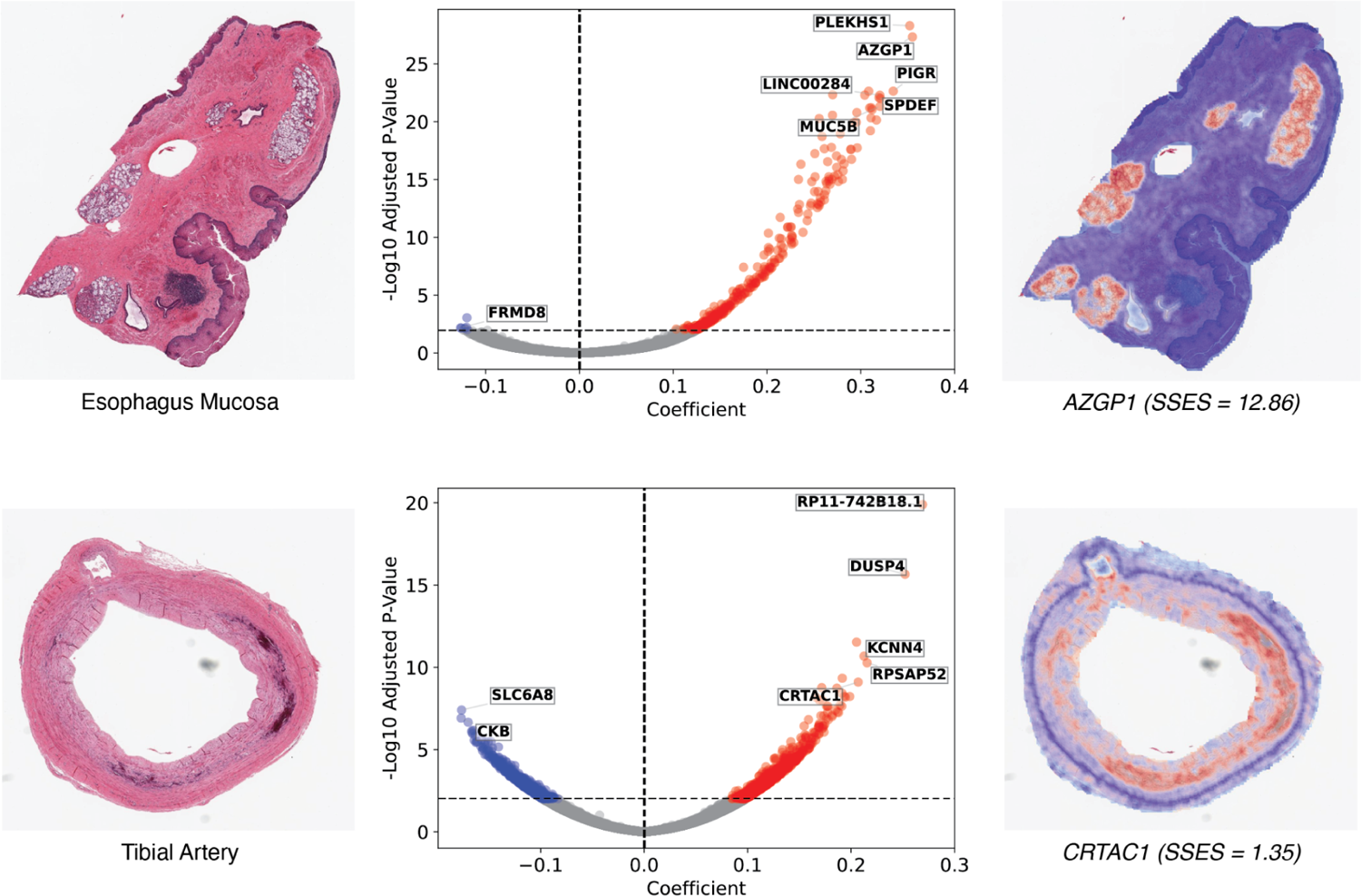
A comparison of differential gene expression analysis and our SSES metric across donors for submucosal gland and calcification proportion. Considering genes with a positive coefficient (up-regulated), we see that our SESS metric when applied to *RNAPath* predictions is able to find genes (e.g. *AZGP1* and *CRTAC1*) that are both significant in DE analysis, but are also highly spatially restricted to submucosal glands and calcification foci, respectively.

In addition to our SSES metric, we computed Moran’s I, a spatial autocorrelation metric commonly used in spatial analysis and more recently in spatial transcriptomics applications^30,31^. A low Moran’s I score indicates a gene is not spatially autocorrelated, with diffuse gene expression across a tissue section, whilst a high Moran’s I score represents high spatial autocorrelation, with expression restricted to specific substructures, tissue neighbourhoods or pathologies. We computed Moran’s I score for all genes across all tissues, using the spatial predictions of *RNAPath*, along with the corresponding patch coordinates (see Data & Code Availability to download full summary statistics). The spatial autocorrelation of genes predicted by *RNAPath* varies significantly, with some genes highly restricted to tissue substructures and others expressed uniformly across the tissue section (Figure 7). Interestingly, as Moran’s I and our spatial predictions are donor specific, we see examples of genes that exhibit subject specific spatial autocorrelation (**Supplementary** Figure 14). This analysis highlights how even without substructure or pathology annotations, using *RNAPath* and derived spatial statistics, it is possible to assign gene expression to specific tissue neighbourhoods.

**Figure 7:**
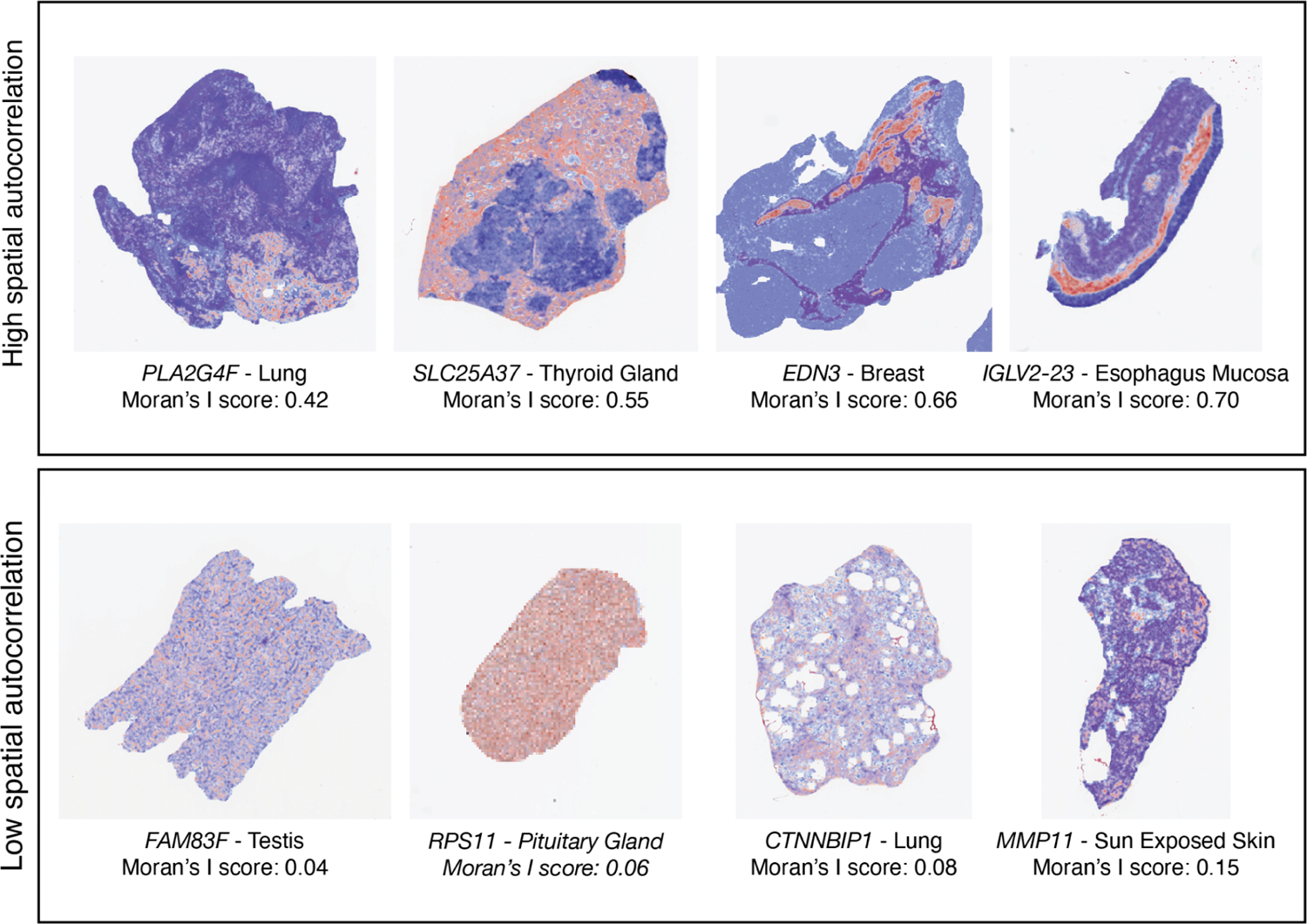
Genes whose spatial autocorrelation is high (top row) and low (bottom row). For example, *IGLV2-23*, a gene whose expression is specific to B-cells, has high spatial autocorrelation and its expression is spatially restricted to regions of focal inflammation below the mucosa. Whereas *RPS11*, a ribosomal subunit, is constitutively expressed across the pituitary gland with low spatial autocorrelation.

## Discussion

Here we use Vision Transformers (ViT) trained using self-distillation with no labels (DINO) to learn histology image representations from 9,068 WSI across 23 healthy human tissues in 838 donors. By doing so, we are able to demonstrate that representations learnt with no labels are able to distinguish tissue substructures and pathological features present in WSI, allowing us to represent each donor’s tissue section as a composite proportion of its underlying tissue substructures and any pathology present. By using these proportions, we show profound inter-tissue variability across donors, demonstrating the detection of unannotated pathologies (e.g. calcification), incorrect target tissue assignment (e.g. esophagus mucosa in muscularis samples), contaminate tissue (e.g. adjacent adipose tissue) and how such variability can inflate eQTL tissue sharing estimates. Additionally, we use such proportions to derive and recapitulate known epidemiological links between breast adiposity and age, as well as novel age, sex and BMI associations. Using our derived tissue substructure and pathological feature proportions we characterise differential gene expression signatures and uncover substructure/pathology genetic associations using GWAS. For example, we find a variant that increases arterial calcification and is associated with coronary angioplasty incidence. Finally, we demonstrate how such proportions can be used to detect interaction eQTLs, in which tissue substructure and pathological variability across donors drive changes in expression in a genotype-dependent manner.

As our histology representations capture intra- and inter-donor variability in tissue morphology, we propose a multiple instance learning model named *RNAPath* that can regress RNA expression levels directly from our learnt histology representations, as well as predict the spatial localisation of a given gene’s expression within a tissue section. We validate our expression predictions by showing substantially better performance of *RNAPath* compared to HE2RNA across a wide range of tissues, and also validate our spatial predictions of known marker genes to ground-truth immunohistochemistry staining. To our knowledge this is the first application of self-supervised learning that allows one to link changes in both pathological and normal histological variability and gene expression variation. Our study advances previous work which was unable to link such histological changes to gene expression variability.^11^

Our work has several limitations and room for future development. First, we work with 128×128 histology image tiles, limiting the resolution of our spatial predictions and segmentations. To overcome this, we extract multiple overlapping tiles to average and smooth our predictions. A natural next step would be to perform cellular or nuclei segmentation and learn self-supervised representations at the single cell-level. We believe this would recapitulate our primary findings but would also allow one to derive more detailed epidemiological, genetic and expression links to specific cell type abundances and their occurrence in pathological and tissue substructures. However, this would be substantially more computationally intensive, as it would require learning and storing millions of cell type representations as compared to thousands of histology tile representations. Second, whilst our discovered GWAS variants recapitulate relationships with known GWAS loci, our genetic analyses are underpowered, both to detect novel GWAS variants and to detect thousands of interaction eQTLs. We believe this will be overcome as larger cohorts with paired histology and genetic data become available, enabling broader discovery but also replication efforts. Third, whilst we demonstrate and validate RNA-expression prediction from histology this will by definition be limited to genes whose variation influences observable morphological differences in tissue sections. To associate histological variation with intracellular gene expression variation beyond morphology would require spatial transcriptomic assays whose current cost does not scale to large numbers of histology sections, with current endeavours profiling only tens of samples^32^. Therefore, we believe there is still significant value in understanding and characterising more deeply, histological and functional genomic associations at the population level. In summary, as histological archives and pathology workflows become digital, we believe there is substantial opportunity for using self-supervised learning to uncover novel, fundamental biology about tissue structure, function and its variability in a population in both healthy and diseased states.

## Methods

### GTEx Cohort description

All analysis is conducted using data from the Genotype Tissue Expression (GTEx) Consortium, which has been described at length in previous publications^8,17,33^. Briefly, GTEx consists of a total of 948 post-mortem donors, in which RNA-seq, Whole Genome Sequencing (WGS), and digitised tissue histology have been collected from up to 54 tissue types. For this study, we utilised GTEx v8 considering the overlap between individual donors who had both RNA-seq and matching tissue histology available and tissues with at least 200 donors genotyped. In total, we utilised n = 9,068 slides, across 23 tissues.

### RNA-seq normalisation

We used normalised TPM values available in the GTEX v8 release. As the GTEx RNA-seq data is not strand-resolved, we only considered lncRNAs that did not overlap with protein coding loci (see Code Availability). Additionally, on a tissue by tissue basis, we considered only genes that were expressed with TPM > 10 in at least 5% of samples. For prediction, we used log normalised TPM values, *log2*(x+1). In total, we considered 21,691 genes.

### GTEx Whole Slide Image histology preprocessing

We downloaded all available Whole Slide Image (WSI) histology data from the GTEx portal. In total, we utilised N=9,068 WSI spanning 23 tissues. WSI were first segmented to separate foreground tissue from background, using a previously published U-net architecture trained on 4,732 H&E slides^34^. Tissue sections were tiled into image patches of fixed dimension (128×128 pixels) and their coordinates stored in hdf5 files. These tiles were used in all downstream analysis.

### Self-supervised features learning from histology patches

After preprocessing, we extracted features from histology tiles giving us a matrix that has as many rows as the number of patches in the WSI and as many columns as the number of features (i.e. 384). To do this, we trained a small vision transformer (ViT-S, output dimension = 384) using DINO^15^, a self-supervised approach on 1.7M histology patches equally sampled across all the 23 tissues from GTEx that we selected for this study. In this self-supervised training regime there are two networks, a student and a teacher model, sharing the same architecture: the student is provided with global and local augmented crops of the input image, whilst the teacher receives only global crops of the same image. Both models output a *k*-dim probability vector (*k* = 65,536) via a temperature softmax along the feature dimension, which can be thought of as a distribution over latent-classes the model is learning to represent. The student-teacher models are trained by minimising the cross entropy (CE) loss between their output distributions:

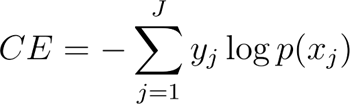

This has the desired effect of encouraging the model to learn local-to-global image correspondences. It can be particularly useful in histology, where cells (local crops) may be specific to much larger tissue structures (global crops). Further details regarding the implementation may be found in the original DINO publication^15^. We modified the augmentation pipeline of DINO’s global and local crops to better capture the relevant features of histology samples, by adding a random modulation of hematoxylin (H) and eosin (E) channels. Images were stain-normalised before being input to the vision transformer to eliminate any effect due to differential stain intensity^35^.

Once obtained, the matrices of tile representations (K × 348, where K is the number of patches of a WSI) for all samples were stored in a HDF5 together with the corresponding upper-left corner coordinates.

### Weakly supervised segmentation of histology images

The segmentation of histology images into regions of interest identifying substructures or pathological features is fundamental to both extract image derived phenotypes (e.g. size of specific tissue regions) and to compute gene enrichments with *RNAPath* predictions.

Our approach is to use weakly supervised clustering: first, we define for each tissue, the substructure and pathologies of interest, considering the pathology notes that are reported together with GTEx samples. Next, we manually annotated, with the support of a clinical histopathologist, a small number of tiles from 5-10 WSI per tissue. To perform these annotations, we used QuPath (v0.4.3)^36^ and a groovy script to produce 128×128 tiles from each annotated area. All annotation tiles are available for download (see Data Availability). For each tile from an annotated class, we perform a forward pass through our trained ViT-S model, obtaining its 384-dim representation. We can then obtain automatic segmentations of the non-annotated WSI by computing the distance between tile representations from unannotated WSI and by using a k-Nearest Neighbours (k = 200) model fitted on the annotated (tissue-specific) dataset to assign classes. We stored the segmentation both as an image and in a dataframe in which the class of each patch is tabulated.

### Differential Expression Analysis of image derived phenotypes

Tissue substructure and pathology proportions were computed by counting the number of patches belonging to each class and normalising by the total number of tissue patches present in the WSI. However, these phenotypes are compositional as the sum across tissue substructure and pathology proportions within a sample equals one, implying that the measured variables are not independent. This dependence may alter the results of downstream statistical analysis. To address this issue, we transformed the compositional values into pivot coordinates^37,38^. Using these pivot transformed proportions, we fit linear models in python adjusting for age, sex, BMI and ischemic time and the first 5 genetic PCs.

### GTEx Whole Genome Sequencing (WGS) quality control

The cohort VCF representing whole genome sequencing variant calls was obtained from dbGaP (accession phs000424.v8.p2). All the analyses described here are based on the GTEx v8 analysis freeze dataset containing 838 individuals and 46,569,704 variants. First, we used somalier^39^ to estimate the ancestry of all samples directly from the cohort VCF. Based on somalier estimations, we then selected only the 699 samples of European (EUR) ancestry based on the 1000G reference populations. Variants were then filtered, retaining only PASS biallelic SNVs. The filtered dataset contained 699 samples and 43,066,451 variants.

To generate a high-quality dataset suitable for GWAS analysis, we further filtered genotypes retaining only those with GQ >= 20 and DP >= 10, and then removed variants with minor allele count < 10, HWE test p-value < 1 × 10^-30^, or missing call rate > 0.05.

The resulting processed dataset containing 11,527,288 variants was converted to PGEN format and used in step 2 of REGENIE for variant association analyses (see below). Variants in this dataset were further processed to generate a set of independent SNVs to be used in step 1 of REGENIE analysis. First, we filtered out variants with HWE test p-value < 1 × 10^-15^, minor allele count < 100, or missing call rate > 0.01, and then we applied LD pruning as implemented by plink2--indep-pairwise method using –indep-pairwise 1000 100 0.5. The final dataset for step 1 included 699 samples and 381,202 variants.

### Genome-wide association analysis (GWAS)

To conduct the genetic association study, we used REGENIE software v3.2.7^40^ with an automated Nextflow pipeline (v1.8.1) (see Data & Code Availability). This pipeline simplifies the testing of multiple phenotypes and automates post-processing steps such as variant clumping for loci identification and annotation of nearby genes. The sample size tested varies depending on the phenotype: 691 donors for tibial artery calcification, 479 for coronary artery calcification and 674 for esophagus mucosa inflammation and vascular congestion. For all four phenotypes we tested autosomal variants adjusting for age, sex, BMI, ischemic time and the first five principal components of genetic ancestry. A minMAC filter of 10 was applied in step 2 of the REGENIE pipeline; and the variants with MAF < 5% were excluded from the final analyses. We considered the standard genome-wide significance threshold (P-value = 5.0 × 10^-8^), but also examined suggestive hits (P-value < 1.0 × 10^-6^). Regional plots, Manhattan plots and quantile-quantile plots were generated with GWASLab (v3.4.21)^41^. All summary statistics are available for download (see Data Availability).

### Interaction eQTL mapping

Interaction eQTLs (ieQTLs) are a specific type of eQTLs that interact with a phenotype (age, sex, environmental factors, etc.) to influence gene expression; as a consequence, ieQTLs can help identify the factors that modulate the effect of genetic variants on gene expression.

To perform interaction eQTL analysis, we used TensorQTL (v1.0.8)^42^, an open source package that allows QTL mapping to be executed on GPUs, resulting in ∼200-300 fold faster computations compared to the CPU-based implementations. Interaction eQTL analysis requires genotypes, gene expression data and an interaction variate (e.g. a phenotype or environmental factor) for each individual. The statistical model is described by the following equation:

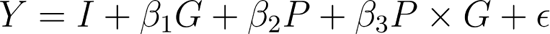

where *I* is the intercept, *G* the genotype, *P* the interaction component (or proxy phenotype), *P × G* represents the interaction term and ε the residual error. We used the same genotype data as per the GWAS (described above). For the gene expression data we utilised the normalised gene expression matrices and covariates provided by GTEx in the cis-eQTLs section of the open access data.

The covariates include the top five genotype components, PEER factors calculated for the expression matrices, sequencing platform, sequencing protocol, sex, age, BMI and ischemic time. As an interaction term, we used the tissue substructure and pathology proportions transformed into pivot coordinates. The sample size varies across tissue type and depends on the number of genotyped donors with both gene expression and WSI of the histology sample available.

### Multiple instance learning applied to gene expression regression

Our novel method, *RNAPath*, follows the multiple instance learning (MIL) paradigm, an approach that enables one to predict patch-level scores whilst having only a sample-level measurement or ground-truth label. *RNAPath* works as follows: Consider a total of WSIs each represented as a bag of image patch embeddings, *X_i_ ∈ R^M×D^* where *M* is the number of image patches for that WSI and *D* is the embedding dimension of each image patch. *M*, termed the bag size in the MIL literature, is variable across WSI as it depends on the size of the tissue section taken. At the level of each WSI, we have *k* regression target variables *y_i_ ∈ R^1×k^*, which are the *log2(x+1)* TPM values for each gene. The model estimates the gene expression at patch level by *G* independent gene-wise linear regressors applied to tile features, where *G* is the number of genes selected for the tissue. We subsequently apply a non-linear activation function (ReLU) to have positive patch-level scores. These scores are then averaged to derive a sample-level prediction; mean squared error (MSE) loss between predicted sample-level expression and bulk RNA-seq is computed to train the model (see Code Availability for full implementation and model architecture).

To train *RNAPath* for each tissue we created a training, validation and test split of 80:10:10, ensuring that tissues from the same individual were present in only one split, to avoid any leakage based on genetic effects shared across tissues. For each sample, we apply dropout both at the bag level (by keeping a random percentage of patches between 70% and 100% of the total number), and at the level of patch representations (p=0.10) to make the training of *RNAPath* more stable and to increase robustness to outliers.

We train *RNAPath* with batch size 1 for a maximum of 200 epochs, using a decaying learning rate scheduler (starting value 1 × 10^-4^); we optimised *RNAPath* using Adam and a Mean Squared Error (MSE) loss function:

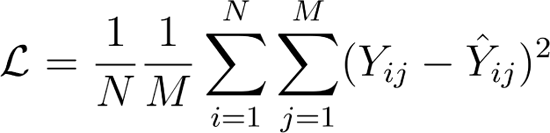

We divided the gene set into groups of size ≤ 500, due to memory restrictions. To limit the time taken by the optimization step, we accumulate the gradients over each of these groups and update weights once all the genes for a sample have been regressed. In total, we trained 23 tissue-specific *RNAPath* models for the regression of gene expression.

As previously described, the patches are extracted from the sample without overlap. However, as an additional form of data augmentation, we compute three additional patch sets (with overlap of 25%, 50% and 75% with respect to the original patch set). By doing so, we achieve four different representations of the same sample, and at each training iteration we can randomly sample one of them. These other patch sets are also useful during inference, as we can average the patch-level logits on the overlapping regions to achieve fine-grained expression heatmaps.

### Substructure Specific Enrichment Score (SSES)

To determine whether a given gene expression prediction was spatially restricted, we devised a substructure specific enrichment score (SSES) that computes a ratio between the mean expression in a given area over the total mean expression, using the patch level predictions:

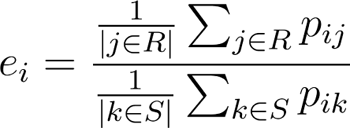

This produces an SSES metric for each gene *i*, *e_i_*, in which *e_i_ > 1* represent genes that are spatially enriched for the given ROI (i.e. the average patch-level expression *p_i_* is higher in the ROI *R* than in the whole sample *S*). Enrichment scores are then averaged across samples, and the final outcome is a matrix reporting the enrichment score for each pair (gene, substructure or localised pathology).

### HE2RNA implementation

*RNAPath* outputs tile-level scores and then averages across tiles for a slide-level aggregated prediction. In our implementation of HE2RNA, we used the author defined model class, available from their published repository, with our 128×128 tiles and self-supervised embeddings. HE2RNA considers bag shapes (number of tiles per WSI) of 8,000 and the number *k* of tiles used in the training step is randomly sampled from the list *L* = [10, 20, 50, 100, 200, 500, 1000, 2000, 5000]. Given that some of our slides have a substantially higher number of tiles, we substituted the absolute numbers in *L* into proportions (*L* / 8000), in order to keep the ratio of tiles used in the training step equal to the original implementation, despite having larger bags. It is worth noting that the original HE2RNA implementation used pre-trained ImageNet feature representations for tiles, which we demonstrate perform significantly worse in representing histological entities. Therefore, this is a conservative comparison in which HE2RNA benefits from using our self-supervised representations.

## Acknowledgements

We would like to thank the Genomics Scientific Support Unit (SSU) at Human Technopole for significant technical and methodological help with this manuscript. The project was supported by a Longevity Impetus Grant from Norn Group.

## Data & Code Availability

GTEx V8 data are accessible via an approved dbGAP application (accession: phs000424.v8.p2). All code, model weights, annotations and summary statistics for this study are publicly available via: https://github.com/GlastonburyC/RNAPath. The nextflow-pipeline used for REGENIE based GWAS is available here: https://github.com/HTGenomeAnalysisUnit/nf-pipeline-regenie.

## References

1. Glastonbury, C. A. et al. Machine Learning based histology phenotyping to investigate the epidemiologic and genetic basis of adipocyte morphology and cardiometabolic traits. PLoS Comput. Biol. 16, e1008044 (2020).

2. Komura, D. et al. Restaining-based annotation for cancer histology segmentation to overcome annotation-related limitations among pathologists. Patterns (N Y) 4, 100688 (2023).

3. Lu, M. Y. et al. Data-efficient and weakly supervised computational pathology on whole-slide images. *Nat*. Biomed. Eng. 5, 555–570 (2021).

4. Fu, Y. et al. Pan-cancer computational histopathology reveals mutations, tumor composition and prognosis. Nat Cancer 1, 800–810 (2020).

5. Deng, J. et al. ImageNet: A large-scale hierarchical image database. in 2009 IEEE Conference on Computer Vision and Pattern Recognition 248–255 (2009).

6. Chen, R. J. & Krishnan, R. G. Self-Supervised Vision Transformers Learn Visual Concepts in Histopathology. arXiv [cs.CV] (2022).

7. Cancer Genome Atlas Research Network et al. The Cancer Genome Atlas Pan-Cancer analysis project. Nat. Genet. 45, 1113–1120 (2013).

8. The GTEx Consortium. The GTEx Consortium atlas of genetic regulatory effects across human tissues. Science 369, 1318–1330 (2020).

9. Schmauch, B. et al. A deep learning model to predict RNA-Seq expression of tumours from whole slide images. Nat. Commun. 11, 3877 (2020).

10. Quiros, A. C. et al. Self-supervised learning in non-small cell lung cancer discovers novel morphological clusters linked to patient outcome and molecular phenotypes. arXiv [cs.CV] (2022).

11. Jones, A., Gundersen, G. W. & Engelhardt, B. E. Linking histology and molecular state across human tissues. bioRxiv 2022.06.10.495669 (2022) doi:10.1101/2022.06.10.495669.

12. Gundersen, G., Dumitrascu, B., Ash, J. T. & Engelhardt, B. E. End-to-end training of deep probabilistic CCA on paired biomedical observations. Proc. Mach. Learn. Res. (2020).

13. Ash, J. T., Darnell, G., Munro, D. & Engelhardt, B. E. Joint analysis of expression levels and histological images identifies genes associated with tissue morphology. Nat. Commun. 12, 1609 (2021).

14. Zhai, X., Kolesnikov, A., Houlsby, N. & Beyer, L. Scaling Vision Transformers. arXiv [cs.CV] (2021).

15. Caron, M. et al. Emerging properties in self-supervised vision transformers. arXiv [cs.CV] 9650–9660 (2021).

16. Dosovitskiy, A., Beyer, L., Kolesnikov, A. & Weissenborn, D. Transformers for image recognition at scale. arXiv preprint arXiv.

17. GTEx Consortium et al. Genetic effects on gene expression across human tissues. Nature 550, 204–213 (2017).

18. Grundberg, E. et al. Mapping cis- and trans-regulatory effects across multiple tissues in twins. Nat. Genet. 44, 1084–1089 (2012).

19. Kim-Hellmuth, S. et al. Cell type-specific genetic regulation of gene expression across human tissues. Science 369, (2020).

20. Costanzo, P. R. et al. Clinical and Etiological Aspects of Gynecomastia in Adult Males: A Multicenter Study. Biomed Res. Int. 2018, 8364824 (2018).

21. Kothari, C., Diorio, C. & Durocher, F. The Importance of Breast Adipose Tissue in Breast Cancer. Int. J. Mol. Sci. 21, (2020).

22. Chen, G. et al. SPDEF is required for mouse pulmonary goblet cell differentiation and regulates a network of genes associated with mucus production. J. Clin. Invest. 119, 2914–2924 (2009).

23. Okuda, K. et al. Localization of Secretory Mucins MUC5AC and MUC5B in Normal/Healthy Human Airways. Am. J. Respir. Crit. Care Med. 199, 715–727 (2019).

24. Duan, L. et al. Novel Susceptibility Loci for Moyamoya Disease Revealed by a Genome-Wide Association Study. Stroke 49, 11–18 (2018).

25. Bai, Y. et al. The intermediate-conductance calcium-activated potassium channel KCa3.1 contributes to alkalinization-induced vascular calcification in vitro. J. Clin. Lab. Anal. 35, e23854 (2021).

26. Glastonbury, C. A. et al. Adiposity-dependent regulatory effects on multi-tissue transcriptomes. Am. J. Hum. Genet. 99, 567–579 (2016).

27. Glastonbury, C. A., Couto Alves, A., El-Sayed Moustafa, J. S. & Small, K. S. Cell-type heterogeneity in adipose tissue is associated with complex traits and reveals disease-relevant cell-specific eQTLs. Am. J. Hum. Genet. 104, 1013–1024 (2019).

28. Donovan, M. K. R., D’Antonio-Chronowska, A., D’Antonio, M. & Frazer, K. A. Cellular deconvolution of GTEx tissues powers discovery of disease and cell-type associated regulatory variants. Nat. Commun. 11, 955 (2020).

29. Digre, A. & Lindskog, C. The Human Protein Atlas-Spatial localization of the human proteome in health and disease. Protein Sci. 30, 218–233 (2021).

30. Qiu, Z. et al. Detection of differentially expressed genes in spatial transcriptomics data by spatial analysis of spatial transcriptomics: A novel method based on spatial statistics. Front. Neurosci. 16, 1086168 (2022).

31. Moran, P. A. P. Notes on continuous stochastic phenomena. Biometrika 37, 17–23 (1950).

32. Hickey, J. W. et al. Organization of the human intestine at single-cell resolution. Nature 619, 572–584 (2023).

33. Carithers, L. J. & Moore, H. M. The genotype-tissue expression (GTEx) project. Biopreservation and biobanking vol. 13 307–308 (2015).

34. Haghighat, M. et al. Automated quality assessment of large digitised histology cohorts by artificial intelligence. Sci. Rep. 12, 5002 (2022).

35. Macenko, M. et al. A method for normalizing histology slides for quantitative analysis. in 2009 IEEE International Symposium on Biomedical Imaging: From Nano to Macro 1107–1110 (2009).

36. Bankhead, P. et al. QuPath: Open source software for digital pathology image analysis. Sci. Rep. 7, 16878 (2017).

37. Hron, K., Filzmoser, P., de Caritat, P., Fišerová, E. & Gardlo, A. Weighted pivot coordinates for compositional data and their application to geochemical mapping. Math. Geosci. 49, 797–814 (2017).

38. Compositional Data Analysis: Theory and Applications. (Wiley-Blackwell, 2011).

39. Pedersen, B. S. et al. Somalier: rapid relatedness estimation for cancer and germline studies using efficient genome sketches. Genome Med. 12, 62 (2020).

40. Mbatchou, J. et al. Computationally efficient whole-genome regression for quantitative and binary traits. Nat. Genet. 53, 1097–1103 (2021).

41. GWASLab: a Python package for processing and visualizing GWAS summary statistics.

42. Taylor-Weiner, A. et al. Scaling computational genomics to millions of individuals with GPUs. Genome Biol. 20, 228 (2019).

