## Supplementary Figures for "Self-supervised learning for characterising histomorphological diversity and spatial RNA expression prediction across 23 human tissue types"

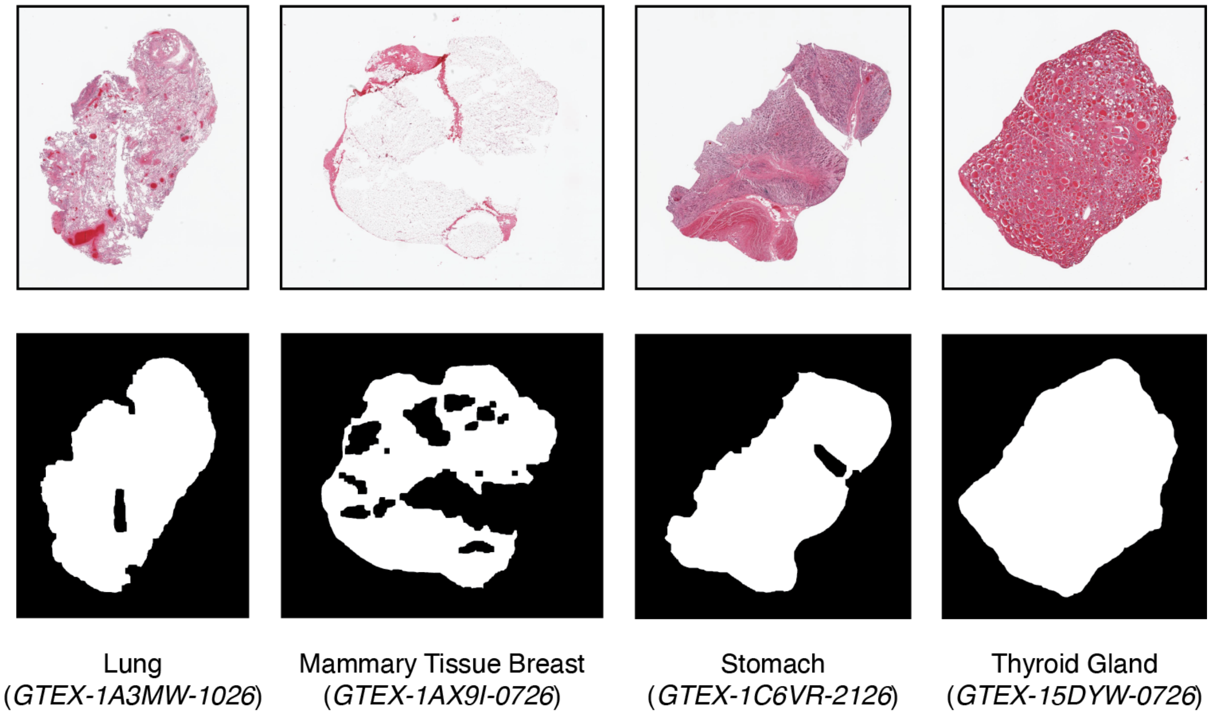


**Supplementary Figure 1**: Binary segmentation of tissue sections of four HE slides from GTEx

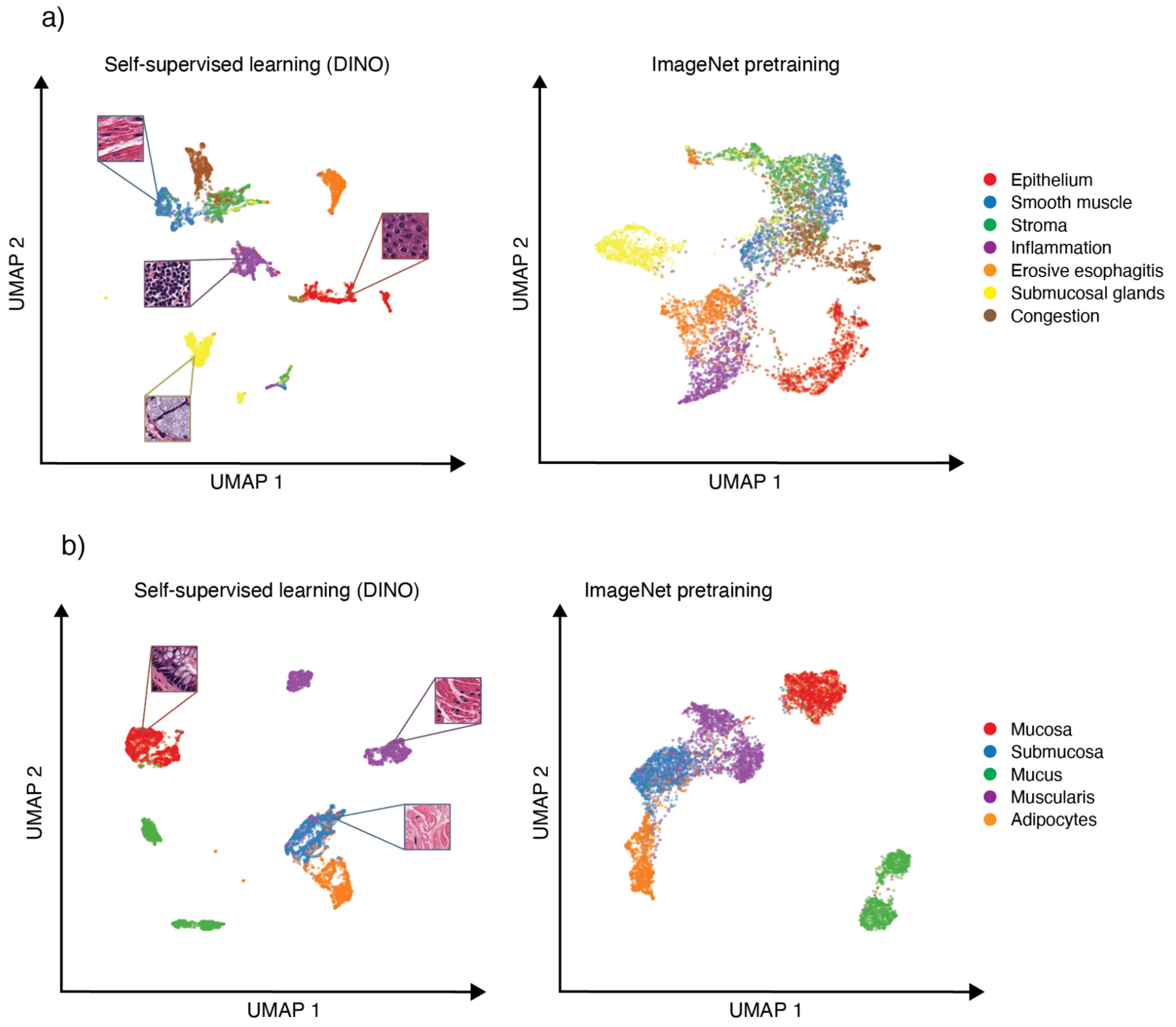


**Supplementary Figure 2**: UMAP embeddings of esophagus mucosa (a) and column (b) patch features from self-supervised ViT-S (trained using DINO from Facebook AI) vs ResNet50 with pretrained weights from ImageNet. Patches have been manually labeled with tissue substructures/pathologies

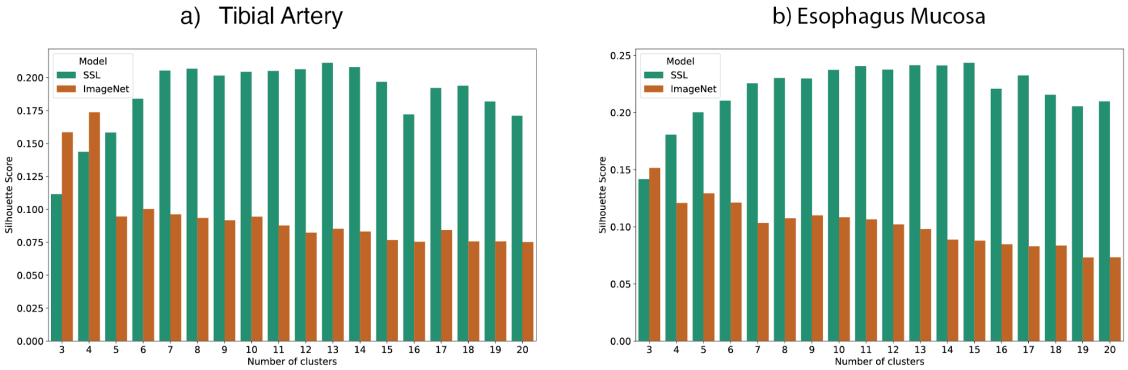


**Supplementary Figure 3**: Comparison of silhouette scores across a several numbers of clusters [3,20]  for tibial artery (a) and esophagus mucosa (b) between pretraining (ImageNet) and self-supervised (SSL) learning of patch features.

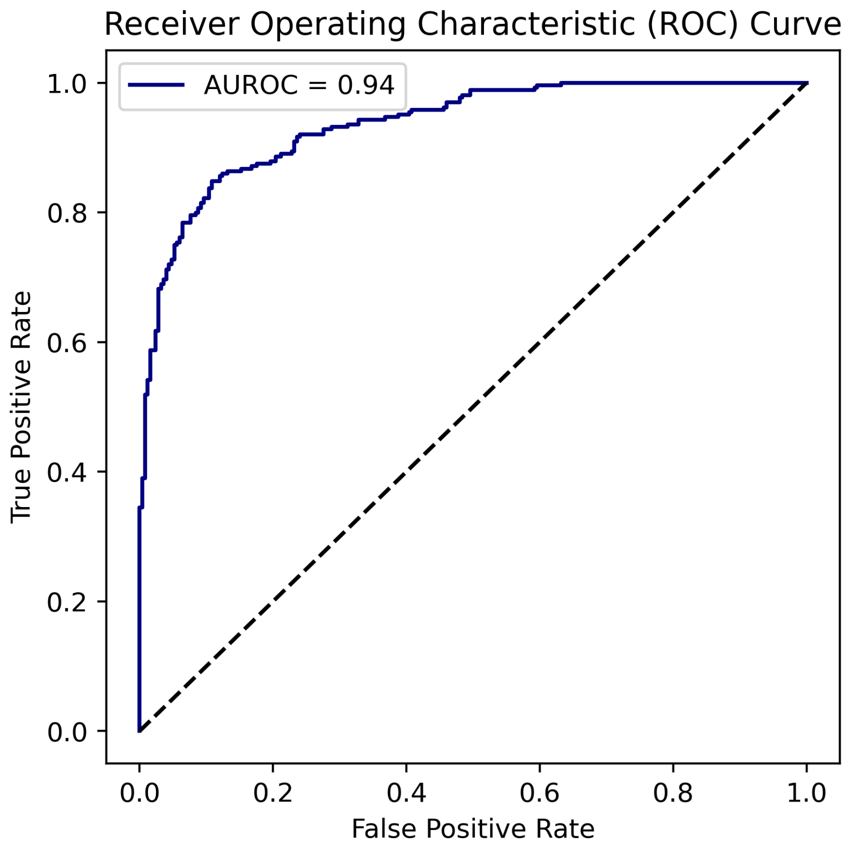


**Supplementary Figure 4:** ROC TPR/FPR pathology notes vs calcification level prediction.

a)

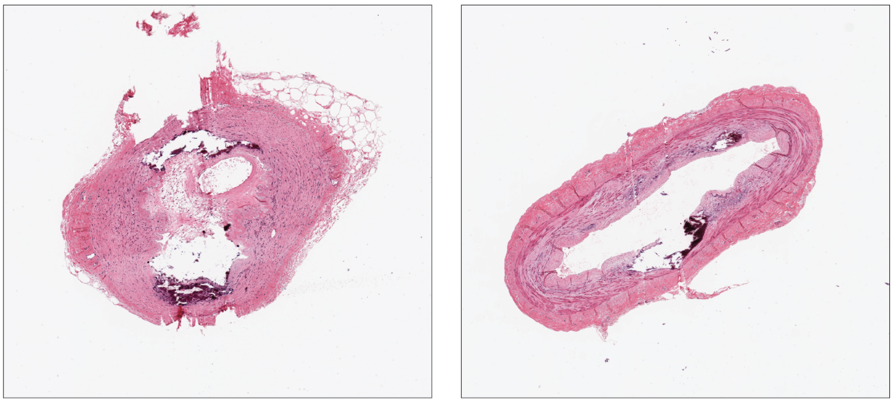

b)

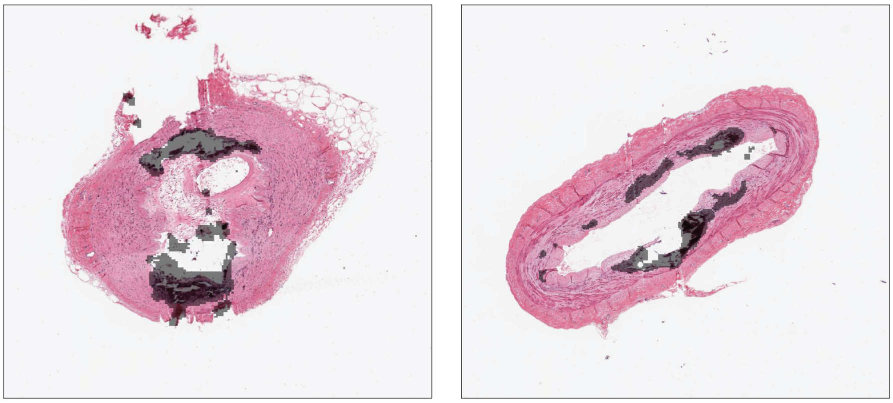

**Supplementary Figure 5**: GTEX-15SKB-0526 and GTEX-X4EP-0826 (a) detected to have tiles containing calcification (b) but were not reported as such in the pathologist notes.

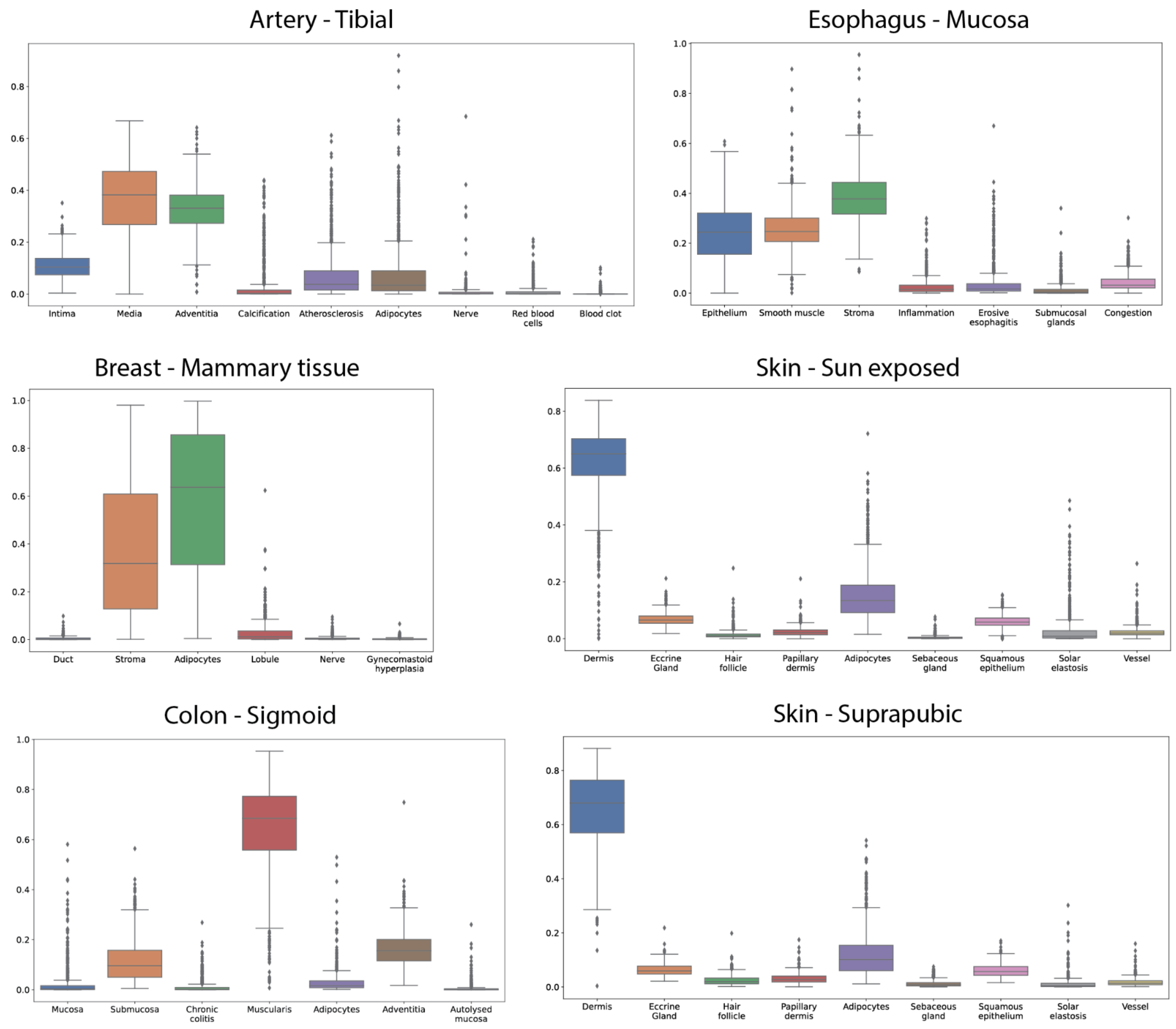


**Supplementary Figure 6**: Variability of derived phenotypes (tissue substructures and localised pathologies) across GTEx donors from 6 example tissues.

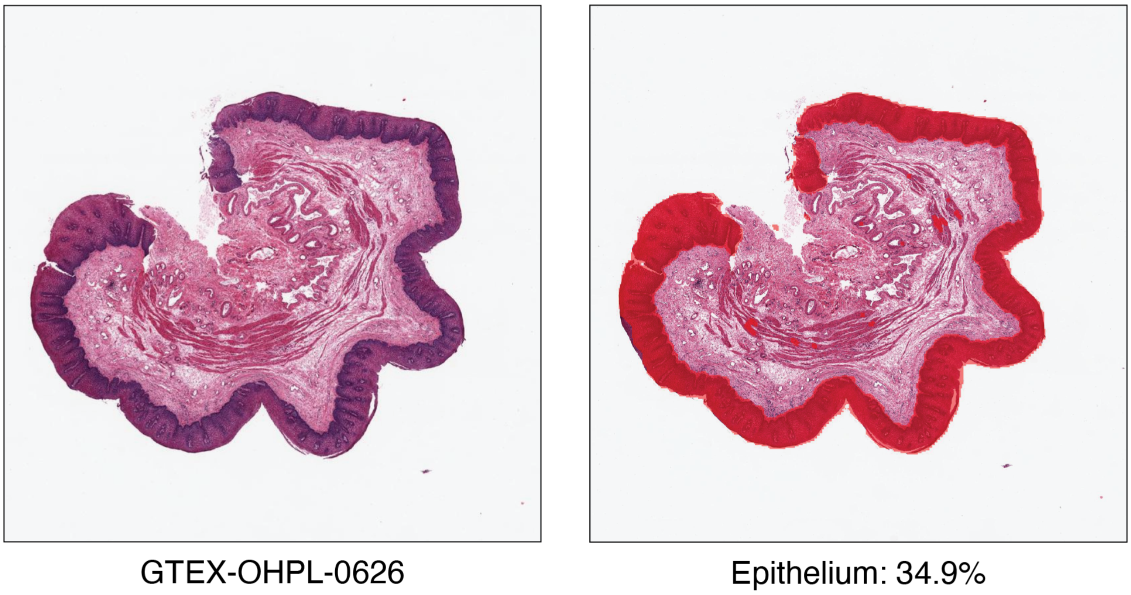


**Supplementary Figure 7:** Esophagus muscularis sample with large mucosal epithelium proportion that has not been trimmed.

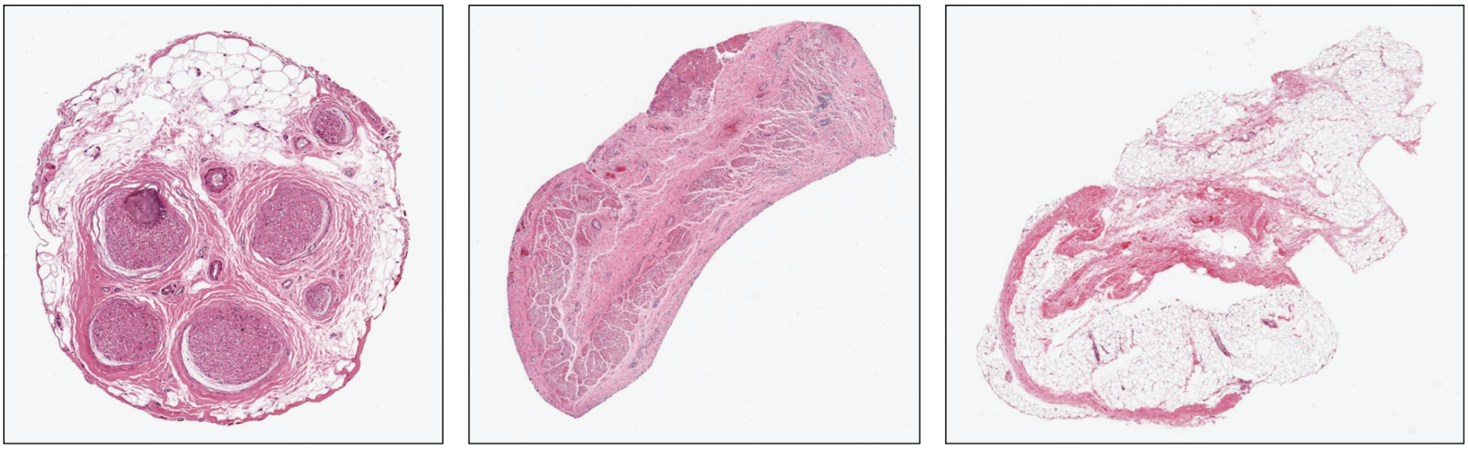

GTEX-P4PQ-1826 (Tibial Artery)
Tunica media: 0.00%

GTEX-1MJK2-1826 (Esophagus Mucosa)
Epithelium: 0.00%

GTEX-15ETS-2526 (Tibial Nerve)
Nerve bundles: 0.00%

**Supplementary Figure 8:** GTEx outliers in the tissue proportions distributions.(Left) Sample stored as tibial artery just having peripheral nerve. (Center) Esophagus mucosa histology without mucosal epithelium. (Right) Tibial nerve sample with no nerve bundle; unknown provenance, not GTEx target according to the pathology notes.

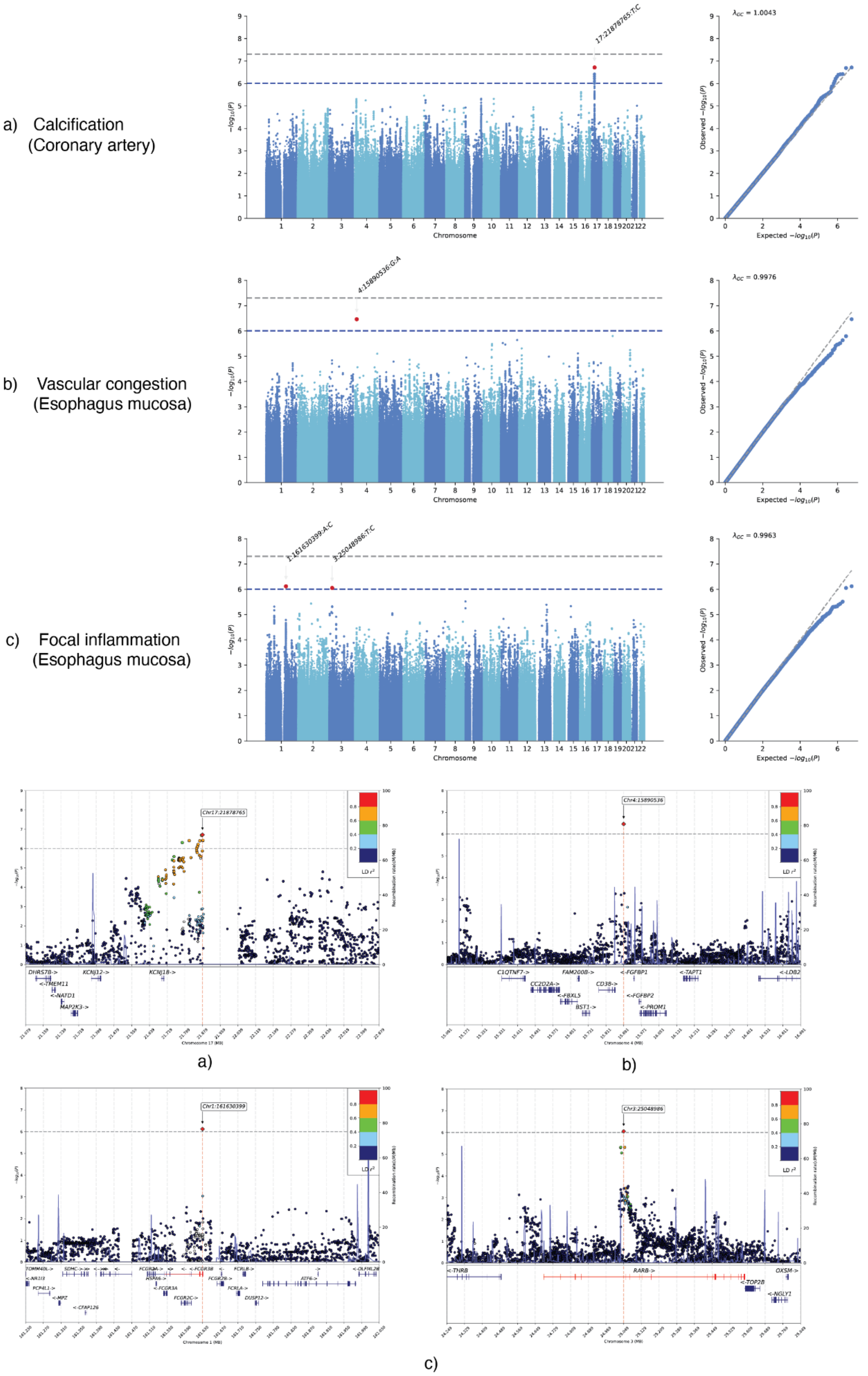


**Supplementary Figure 9:** Manhattan, QQ and locus zoom plots for four variants associated with calcification, vascular congestion and focal inflammation.

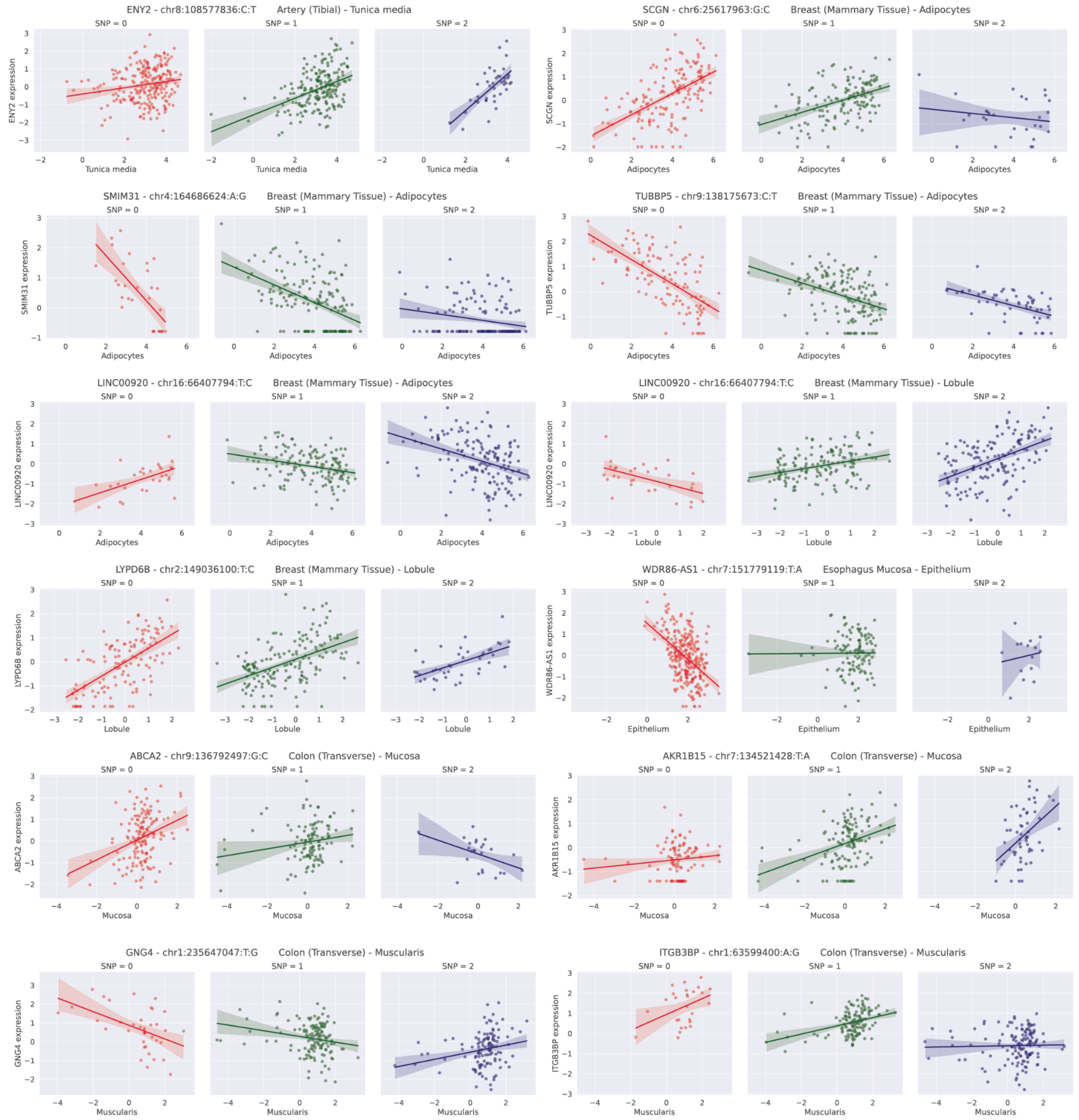


**Supplementary Figure 10:** Example interaction eQTL pivot plots for several tissue substructures and pathologies.

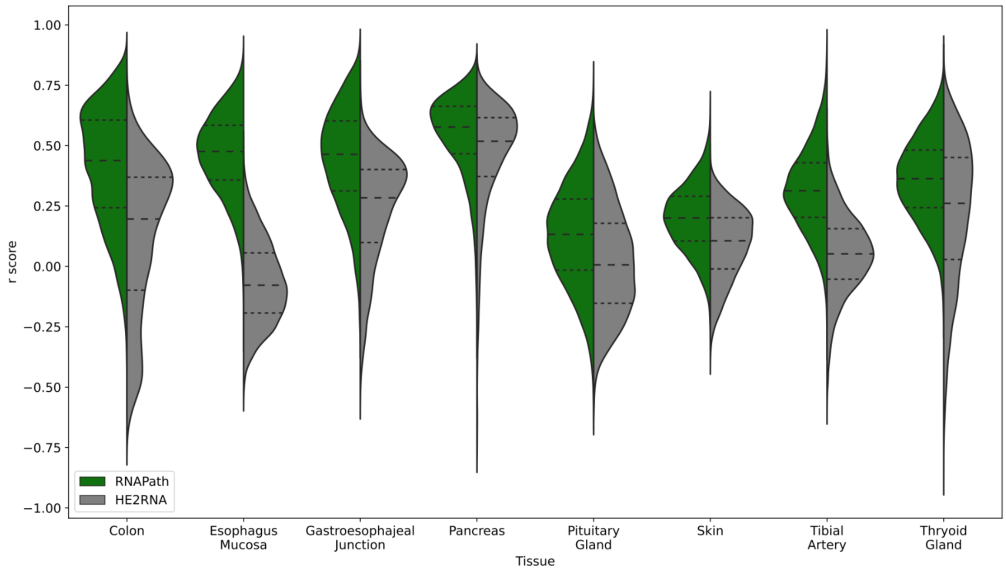


**Supplementary Figure 11:** Comparison of RNA-expression prediction accuracy by RNAPath as compared to HE2RNA across 8 example tissues.

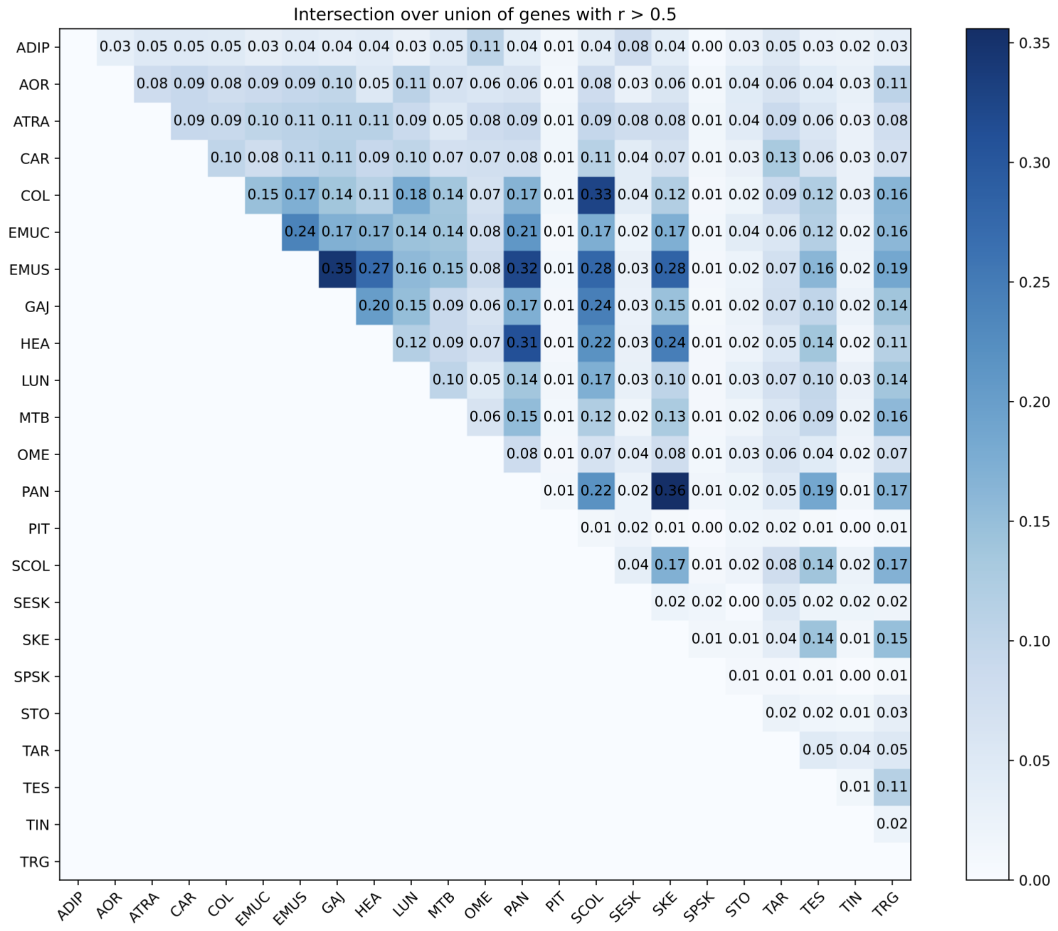


**Supplementary Figure 12:** Intersection over union (IoU) of genes regressed with correlation r-score > 0.5. The couples of tissues sharing most of well-predicted genes are pancreas with skeletal muscle, esophagus muscularis with gastroesophageal junction and transverse colon with sigmoid colon.

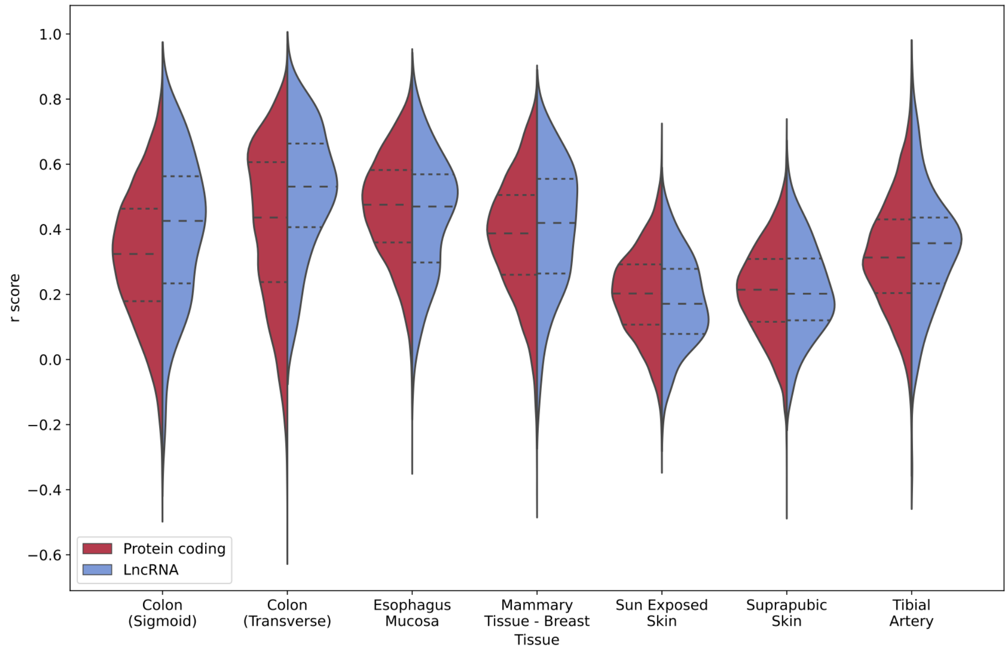


**Supplementary Figure 13:** Comparison of RNA-expression prediction accuracy between protein coding genes and long non-coding RNAs across the 7 annotated tissues.

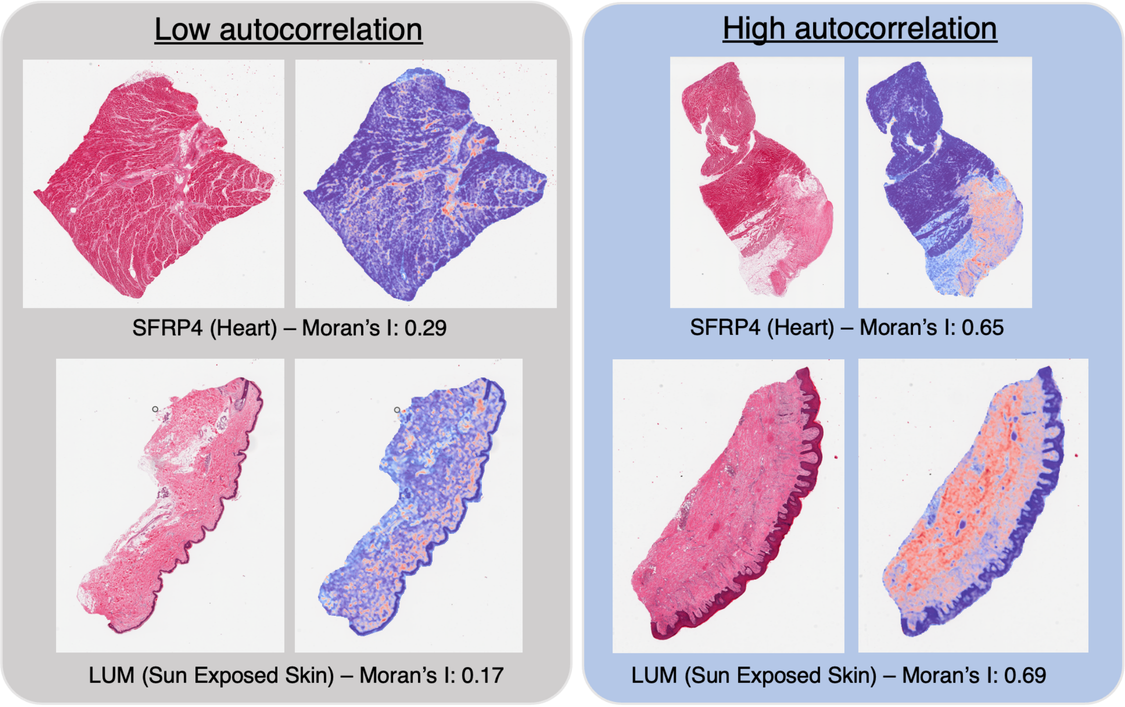


**Supplementary Figure 14:** Examples of genes across donors which exhibit donor-specific spatial autocorrelation. For example, *SFRP4*
