## Supplementary Tables for "Self-supervised learning for characterising histomorphological diversity and spatial RNA expression prediction across 23 human tissue types"

| Tissue | Phenotype | Annotated Tiles | Accuracy |
|---|---|---|---|
| *Artery (Tibial)* | Tunica intima | 1,038 | 0.92 |
| | Tunica media | 5,365 | 0.89 |
| | Tunica adventitia | 3,172 | 0.91 |
| | Calcification | 1,058 | 0.95 |
| | Atherosclerosis | 1,780 | 0.94 |
| | Adipocytes | 5,297 | 0.97 |
| | Nerve | 1,376 | 0.98 |
| | Red blood cells | 465 | 0.96 |
| | Blood clot | 228 | 0.90 |
| *Colon (Transverse)* | Mucosa | 2,680 | 0.97 |
| | Submucosa | 4,030 | 0.97 |
| | Mucus | 1,215 | 0.99 |
| | Muscularis | 2,752 | 0.94 |
| | Adipocytes | 953 | 0.92 |
| *Colon (Sigmoid)* | Mucosa | 2,692 | 0.94 |
| | Submucosa | 2,628 | 0.73 |
| | Chronic colitis | 541 | 0.92 |
| | Muscularis | 5,545 | 0.97 |
| | Adipocytes | 823 | 0.86 |
| | Adventitia | 1,030 | 0.83 |
| | Autolysed mucosa | 196 | 0.82 |
| *Esophagus Mucosa* | Epithelium | 1,530 | 0.92 |
| | Smooth muscle | 2,963 | 0.87 |
| | Stroma | 4,328 | 0.84 |
| | Inflammation | 884 | 0.96 |
| | Erosive esophagitis | 747 | 0.98 |
| | Submucosal gland | 1,822 | 0.90 |
| | Congestion | 630 | 0.87 |
| *Mammary Tissue Breast* | Duct | 471 | 0.47 |
| | Stroma | 11,399 | 0.97 |
| | Adipocytes | 12,820 | 0.98 |
| | Lobule | 803 | 0.93 |
| | Nerve | 328 | 0.80 |
| | Gynecomastoid hyperplasia | 262 | 0.80 |
| *Skin* | Dermis | 1,464 | 0.98 |
| | Eccrine gland | 763 | 0.88 |
| | Hair follicle | 960 | 0.86 |
| | Papillary dermis | 218 | 0.25 |
| | Adipocytes | 2,727 | 0.92 |
| | Sebaceous gland | 653 | 0.84 |
| | Squamous epithelium | 470 | 0.83 |
| | Solar elastosis | 178 | 0.87 |

| | | | |
|---|---|---|---|
| | Vessel | 630 | 0.82 |

**Supplementary Table 1**: Accuracy of all the annotated image derived phenotypes across 6 tissues.

| Tissue | Phenotype | Mean | Standard Deviation | Min | Max |
|---|---|---|---|---|---|
| *Artery (Tibial)* | Tunica intima | 0.11 | 0.05 | 0.00 | 0.35 |
| | Tunica media | 0.36 | 0.14 | 0.00 | 0.67 |
| | Tunica adventitia | 0.33 | 0.09 | 0.01 | 0.64 |
| | Calcification | 0.03 | 0.07 | 0.00 | 0.44 |
| | Atherosclerosis | 0.07 | 0.09 | 0.00 | 0.61 |
| | Adipocytes | 0.08 | 0.11 | 0.00 | 0.92 |
| | Nerve | 0.01 | 0.03 | 0.00 | 0.68 |
| | Red blood cells | 0.01 | 0.02 | 0.00 | 0.21 |
| | Blood clot | 0.00 | 0.01 | 0.00 | 0.10 |
| *Colon (Transverse)* | Mucosa | 0.16 | 0.11 | 0.00 | 0.82 |
| | Submucosa | 0.35 | 0.09 | 0.03 | 0.84 |
| | Mucus | 0.07 | 0.07 | 0.00 | 0.21 |
| | Muscularis | 0.33 | 0.14 | 0.00 | 0.25 |
| | Adipocytes | 0.10 | 0.08 | 0.00 | 0.72 |
| *Colon (Sigmoid)* | Mucosa | 0.03 | 0.06 | 0.00 | 0.58 |
| | Submucosa | 0.12 | 0.09 | 0.01 | 0.56 |
| | Chronic colitis | 0.01 | 0.03 | 0.00 | 0.27 |
| | Muscularis | 0.64 | 0.18 | 0.01 | 0.95 |
| | Adipocytes | 0.03 | 0.05 | 0.00 | 0.53 |
| | Adventitia | 0.16 | 0.07 | 0.02 | 0.75 |
| | Autolysed mucosa | 0.01 | 0.02 | 0.00 | 0.26 |
| *Esophagus Mucosa* | Epithelium | 0.23 | 0.13 | 0.00 | 0.61 |
| | Smooth muscle | 0.26 | 0.09 | 0.00 | 0.90 |
| | Stroma | 0.38 | 0.11 | 0.00 | 0.96 |
| | Inflammation | 0.03 | 0.03 | 0.00 | 0.30 |
| | Erosive esophagitis | 0.04 | 0.06 | 0.00 | 0.67 |
| | Submucosal gland | 0.01 | 0.03 | 0.00 | 0.34 |
| | Congestion | 0.05 | 0.04 | 0.00 | 0.30 |
| *Mammary Tissue Breast* | Duct | 0.00 | 0.01 | 0.00 | 0.10 |
| | Stroma | 0.38 | 0.28 | 0.00 | 0.98 |
| | Adipocytes | 0.58 | 0.31 | 0.00 | 1.00 |
| | Lobule | 0.03 | 0.05 | 0.00 | 0.62 |
| | Nerve | 0.01 | 0.01 | 0.00 | 0.09 |
| | Gynecomastoid hyperplasia | 0.00 | 0.00 | 0.00 | 0.07 |
| *Skin* | Dermis | 0.62 | 0.13 | 0.00 | 0.84 |
| | Eccrine gland | 0.07 | 0.02 | 0.02 | 0.21 |
| | Hair follicle | 0.01 | 0.02 | 0.00 | 0.25 |
| | Papillary dermis | 0.02 | 0.02 | 0.00 | 0.21 |

| | | | | | |
|---|---|---|---|---|---|
| *(Sun exposed)* | Adipocytes | 0.15 | 0.09 | 0.02 | 0.72 |
| | Sebaceous gland | 0.00 | 0.01 | 0.00 | 0.08 |
| | Squamous epithelium | 0.06 | 0.02 | 0.00 | 0.15 |
| | Solar elastosis | 0.03 | 0.05 | 0.00 | 0.49 |
| | Vessel | 0.02 | 0.02 | 0.00 | 0.26 |
| *Skin (Suprapubic)* | Dermis | 0.65 | 0.14 | 0.00 | 0.88 |
| | Eccrine gland | 0.06 | 0.02 | 0.02 | 0.22 |
| | Hair follicle | 0.02 | 0.02 | 0.00 | 0.20 |
| | Papillary dermis | 0.03 | 0.02 | 0.00 | 0.17 |
| | Adipocytes | 0.12 | 0.09 | 0.01 | 0.54 |
| | Sebaceous gland | 0.01 | 0.01 | 0.00 | 0.07 |
| | Squamous epithelium | 0.06 | 0.02 | 0.02 | 0.17 |
| | Solar elastosis | 0.01 | 0.02 | 0.00 | 0.30 |
| | Vessel | 0.02 | 0.01 | 0.00 | 0.16 |
| *Coronary Artery* | Calcification | 0.03 | 0.07 | 0.00 | 0.57 |
| | Adipocytes | 0.34 | 0.22 | 0.00 | 0.95 |
| *Tibial Nerve* | Adipocytes | 0.29 | 0.14 | 0.01 | 0.97 |
| | Nerve | 0.49 | 0.14 | 0.00 | 0.88 |
| *Esophagus Muscularis* | Epithelium | 0.00 | 0.02 | 0.00 | 0.35 |
| | Inflammation | 0.00 | 0.01 | 0.00 | 0.13 |
| | Glands | 0.00 | 0.00 | 0.00 | 0.08 |

**Supplementary Table 2**: Mean, standard deviation, minimum and maximum value of image derived phenotypes.

| Tissue | Median r score | #Genes r > 0.5 | #Genes r > 0.8 | #Genes regressed |
|---|---|---|---|---|
| *Adipose Tissue* | 0.25 | 946 | 2 | 11,413 |
| *Aorta* | 0.22 | 629 | 9 | 11,591 |
| *Atrial Appendage* | 0.40 | 2,195 | 58 | 8,857 |
| *Colon (Transverse)* | 0.44 | 4,948 | 590 | 12,002 |
| *Colon (Sigmoid)* | 0.33 | 2,356 | 127 | 11,683 |
| *Coronary Artery* | 0.19 | 1,517 | 136 | 12,163 |
| *Esophagus Mucosa* | 0.48 | 4,869 | 278 | 10,975 |
| *Esophagus Muscularis* | 0.35 | 2.598 | 123 | 11,014 |
| *Gastroesophageal Junction* | 0.46 | 4,866 | 639 | 11,217 |
| *Heart* | 0.65 | 6,097 | 1,384 | 7,538 |
| *Lung* | 0.35 | 2,983 | 224 | 13,115 |
| *Mammary Tissue Breast* | 0.39 | 3,327 | 129 | 12,471 |
| *Omentum* | 0.37 | 2,067 | 4 | 11,582 |
| *Pancreas* | 0.58 | 5,042 | 378 | 7,370 |
| *Pituitary Gland* | 0.13 | 475 | 3 | 12,788 |
| *Skeletal Muscle* | 0.52 | 4,516 | 289 | 8,305 |

| | | | | |
|---|---|---|---|---|
| *Skin (Sun Exposed)* | 0.20 | 138 | 0 | 11,538 |
| *Skin (Suprapubic)* | 0.21 | 187 | 0 | 11,514 |
| *Stomach* | 0.39 | 3,174 | 36 | 11,530 |
| *Testis* | 0.50 | 7,473 | 960 | 14,994 |
| *Thyroid Gland* | 0.36 | 2,817 | 49 | 12,970 |
| *Tibial Artery* | 0.31 | 1,648 | 73 | 11,282 |
| *Tibial Nerve* | 0.19 | 298 | 5 | 12,614 |

**Supplementary Table 3**: Summary of RNAPath results across tissues (median correlation, number of genes with correlation coefficient $r > 0.5$ and $r > 0.8$, total number of regressed genes).
